# Time-resolved cryo-EM using a combination of droplet microfluidics with on-demand jetting

**DOI:** 10.1101/2022.10.21.513149

**Authors:** Stefania Torino, Mugdha Dhurandhar, Annelore Stroobants, Raf Claessens, Rouslan G. Efremov

## Abstract

Using single particle cryogenic electron microscopy (cryo-EM) high-resolution structures of proteins in different conformations can be reconstructed. Protein function often involves transient functional conformations, which can be resolved using time-resolved cryo-EM (trEM). In trEM, reactions are arrested after a defined delay time by rapid vitrification of protein solution on the EM grid. Despite the increasing interest in trEM among the cryo-EM community, making trEM samples with a time resolution below 100 ms remains challenging. Here we report the design and the realization of a time-resolved cryo-plunger that combines a droplet-based microfluidic mixer with a laser-induced generator of microjets that allows rapid initiation of reaction and rapid plunge-freezing of cryo-EM grids. Using this approach, a time resolution of 5 ms was achieved and the protein density map was reconstructed to a spatial resolution of 2.1 Å. We performed trEM experiments on GroEL:GroES chaperonin complex, these resolved the kinetics of the complex formation and visualized putative short-lived conformations of GroEL-ATP complex.

## Introduction

Proteins sample a complex conformational landscape shaped by environment, ligand binding, protein-protein interactions, and charge exchange^1^. The shape of the landscape determines how the stochastic walk through the conformations is converted into directed motions^2^, how chemical reactions are coupled to useful work^3^, how interaction partners are selected^4,5^, and it also defines the lifetimes of functionally active states^6–8^. Reconstructing the conformational landscape experimentally, with its depressions representing stable and metastable states, and barriers defining kinetics and direction of conformational transitions remains challenging^9^.

Single particle cryogenic electron microscopy (cryo-EM) resolves conformational states of proteins frozen in vitreous ice with high spatial resolution^10–14^. Conventional preparation of cryo-EM grids by blotting techniques^15^ requires several seconds from sample application on the grid until plunge-freezing. As the result, many functionally important transient conformations with short lifetimes do not populate even under continuous turnover conditions^16,17,18^, and therefore are not accessible to traditional cryo-EM.

Using time-resolved cryo-EM (trEM) both the kinetics of the conformational transitions and the high-resolution structures of the transient states can in principle be resolved^11,19,20^. In trEM, the reaction is initiated by mixing^21–25^ or flash photolysis^26,27^ and is arrested by plunge freezing at sequential time points as the system relaxes towards its kinetic equilibrium^28,29^. Because of the rapid cooling during vitrification, trEM has the potential for reaching microsecond time resolution^15^.

trEM was pioneered in 1990^th^. First, flash-photolysis time-resolved plungers were built^26,27^, followed by mixing-based plungers^21^. These methods were applied to study the transient conformations in 2D crystals of bacteriorhodopsin^19,26^ and the open conformation of acetylcholine receptor^30^. Later, microfluidic mixers coupled to gas-assisted atomizer^24,31^, or electrospray^25^, were introduced to study the dynamics of ribosome^11^ and actin-myosin interactions^25^. More recently, on-grid mixing by overlaying microdroplets produced by a drop-on-demand (DOD) piezoelectric dispenser was also used to perform trEM experiments with a time resolution of around 100 ms^32^.

Despite a 30-year history of methodological developments, applications of this high-potential technique remained elusive due to excessive sample consumption, poor reproducibility of EM grids^23–25,33^, or limited time resolution^32^.

To render millisecond trEM feasible and applicable to protein samples available in small amounts, we developed an instrument that combines a rapid low-consumption mixer with a DOD spray generator. We show that trEM samples can be prepared with a time resolution of 5 ms while consuming less than 100 nL of protein solution per EM grid, making our trEM sample preparation potentially applicable to scarce proteins while maintaining high time resolution. The prepared trEM grids display complete mixing of large proteins and are suitable for high-resolution imaging. Furthermore, benchmarking the device with molecular machine chaperonin GroEL:GroES demonstrated that near-atomic resolution reconstructions can be routinely obtained with reaction time down to 10 ms resolving kinetics of complex assemblies and visualizing transient conformational states.

## Results

### Rational and characteristics of the time-resolved plunger

To solve the bottleneck of trEM sample preparation, that is to achieve the reproducible generation of sprayed microdroplets with short mixing times while maintaining low sample consumption, we thought of designing a microfluidic device that combines a fast mixer with a controlled and tunable generator of microdroplet spray and a rapid plunger. After considering and testing several mixing and spray-generating methods, we designed a microfluidic chip that uses a droplet-microfluidics-based mixer^34^ and a DOD spray generator^35^. The practical realization of this approach resulted in the microfluidic chip and the integrated plunger device shown in **Fig. 1a, b,** and **Supplementary Fig. 1**.

**Figure 1.**
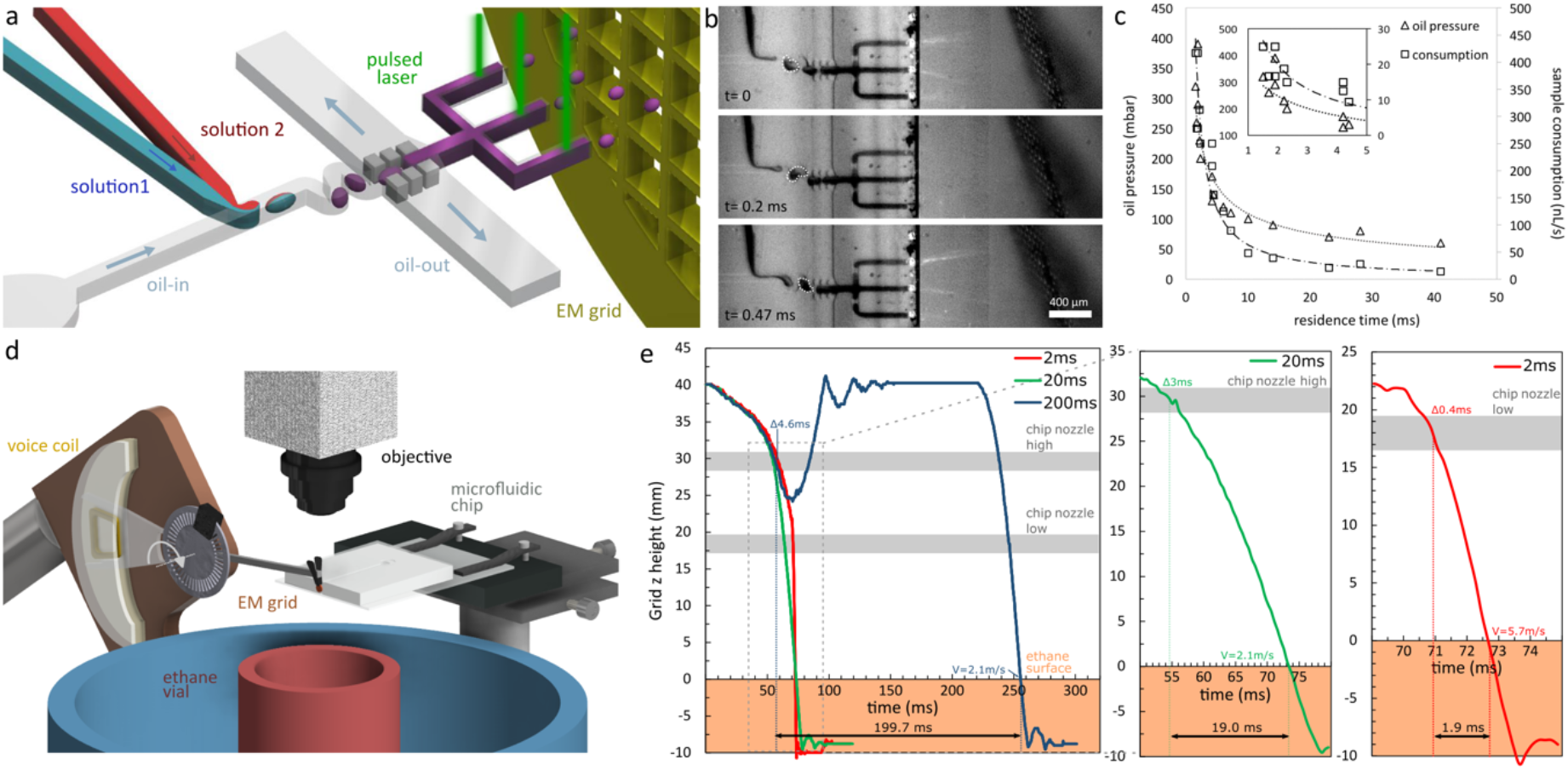
Design and characteristics of the time-resolved plunger. **a**, Schematics, and operation principle of the microfluidic chip that comprises a droplet generator, a serpentine that enhances solutes mixing inside the droplets, a droplet merger, and nozzles where protein solution jets are generated by LIC.**b**, Snapshots of the chip operation. Selected frames from a high-speed video (15000 fps) show the movement of a droplet through the serpentine channel and spray ejection toward the grid moving in front of the nozzles. The residence time was 4.5 ms. **c**, Dependence of residence time of the aqueous solutes (from the initiation of mixing until ejection) on pressure-driving oil flow. The ratio between the oil pressure and the aqueous phase varied in a range between 1 and 2. **d**, 3D drawing of the plunger. Functionally important elements are labeled. **e**, Traces of plunger arm Z-coordinate are shown for three plunge times. The arm is moved slowly close to the nozzle where it is accelerated and decelerated before entering the liquid ethane vial. The time of grid passage before nozzles and plunge time (t_plunge_) are indicated. The grey band shows an area where the grid faces the nozzles, and the orange-filled area represents liquid ethane.

In the microfluidic chip (**Fig. 1a**) two aqueous solutions are combined and encapsulated into microdroplets in the oil phase with a volume of about 100 pL. The content of the droplets is mixed by chaotic advection which is enhanced by passing the droplets through a winding channel^34^. We found that mixing efficiency exceeds 50 % in a serpentine consisting of three turns (**Supplementary Fig. 2**). The mixing was completed downstream of the chip upon droplet merging and spray generation^36^ (see below). To minimize the amount of oil sprayed on the EM grids the aqueous droplets were merged into a continuous stream using a pillar-induced droplet merger module^37,38^ whereby the aqueous and the oil phases are separated into different channels (**Fig. 1a, b, Supplementary Fig. 1c**).

The mixed aqueous solution is directed into the nozzle where airborne droplets are generated by laser-induced cavitation (LIC)^39,40^. A pulsed laser focused on a microfluidic channel next to the nozzle is absorbed by a dye dissolved in the aqueous phase generating a bubble with a microsecond lifetime. The bubble expansion and collapse result in a rapid oscillation of the meniscus and jetting of the liquid^39^ (**Supplementary Video 1**). LIC has the advantage of local actuation tunable by varying the laser power and frequency while preserving the simplicity of the microfluidic chip design. We found that a pulsed Q-switched Nd:YAG laser (WL 532 nm, 6 ns) at laser power per pulse and per focal point of 4-10 μJ generated a stable spray when amaranth dye was used at concentrations above 8 mM. The velocity of generated droplets was in the range between 3 and 6 m/s (**Supplementary Fig. 3**).

The number of droplets deposited on each EM grid and consequently the number of areas suitable for high-resolution cryo-EM imaging were multiplexed using three parallel nozzles and multiple laser focal points obtained using a diffractive beam splitter (**Fig. 1a, b, Supplementary Fig. 1**). The jets were ejected towards the EM grid moving in a vertical plane in front of the nozzle (**Fig. 1d, Supplementary Video 1**).

The residence time of the aqueous solution in the microfluidic chip, the timespan from droplet formation until droplet ejection (tchip), was tunable between 1 and 40 ms (**Fig. 1c**) at flow-driving pressure between 50 and 150 mbar for aqueous phase and between 50 and 400 mbar for the oil phase. The corresponding flow rates of the aqueous phase varied between ~20 and 420 nL/s per channel (**Fig. 1c**). A somewhat counterintuitive non-linear relationship between the applied pressure and flow rate (**Fig. 1c**) is a well-known phenomenon in droplet systems, wherein the linearity is lost due to the discontinuous nature of the flow^41^.

Despite the apparent invasiveness of LIC, its influence on protein integrity is limited. An estimated volume fraction below 3 % per each ejected droplet is exposed to the direct laser beam (**Supplementary Video 1**), whereas the residual heat wave is localized within radius of 15 μm, short-lived (~1 ms), and rises temperature by around 10 ^o^C^42^.

The effect of LIC on protein integrity was assessed experimentally. We measured the catalytic activity of ß-galactosidase before and after passing through the microfluidic chip and being ejected by LIC (**Supplementary Fig. 4**). More than 80 % of the protein activity was preserved. A continuous activity reduction from 99±1 % to 81±2 % was observed as the residence time decreased from 43 to 3 ms. It was likely caused by decrease in the volume of the ejected microjets and can likely be mitigated through a more accurate control of the liquid flow.

The reaction delay-time (t_reaction_), the timespan from the initiation of mixing until the grid plunge-freezing, is a sum of three terms:

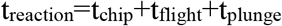

where, t_flight_ is the time of flight between the chip nozzles and the EM grid, and t_plunge_ is the time of grid transfer between the chip nozzle and liquid ethane. In our setup, the EM grid passes at a distance from the nozzle of around 0.5 mm (**Fig 1b, d**, **Supplementary Video 1**), and the t_flight_ is estimated to be around 0.5 ms. Therefore, the reaction time is determined by the combination of longer residence and plunge times.

A custom miniature plunger was built to fit into the constrained space of the microfluidic and optical setup (**Fig. 1b, Supplementary Fig. 1a, b**). EM grid is mounted on an arm driven by a rotational voice coil which accelerates and decelerates the grid at up to 10^4^ m/s^2^ while maintaining accurate control over the arm position (**Fig. 1e, Supplementary Video 1**). The plunge time is programmatically set to any value above the minimum time of 2 ms. For a t_plunge_ exceeding 100 ms, the plunger arm after spray deposition is lifted to the starting position in a local high-humidity environment to minimize the sample evaporation (**Fig. 1e,** 200 ms trace). The optimized profile of the arm acceleration maximizes the number of deposited microjets during the passage of the grid in front of the nozzle. The time of spraying on the grid varies between 4.6 ms, for plunge times above 80 ms, and 0.4 ms, for the shortest plunge time of 2 ms.

The sample consumption was minimized using a pulsed application of fluid flow synchronized with the movement of the plunger arm (**Supplementary Video 1**). The pressure-controlled flow was actuated for a duration of around 200 ms which reduced sample consumption per grid to below 100 nL. In practice, tens of cryo-EM grids were routinely prepared using 20 μL of protein solution.

### Benchmark with test proteins

The time-resolved plunger was benchmarked using common test proteins: mouse heavy chain apoferritin (MW 487 kDa) and *Escherichia coli* β-galactosidase (MW 464 kDa). The proteins were supplied into separate channels of the microfluidic chip and plunged with reaction delay times of 5, 35, and 205 ms (**Fig. 2**).

**Figure 2.**
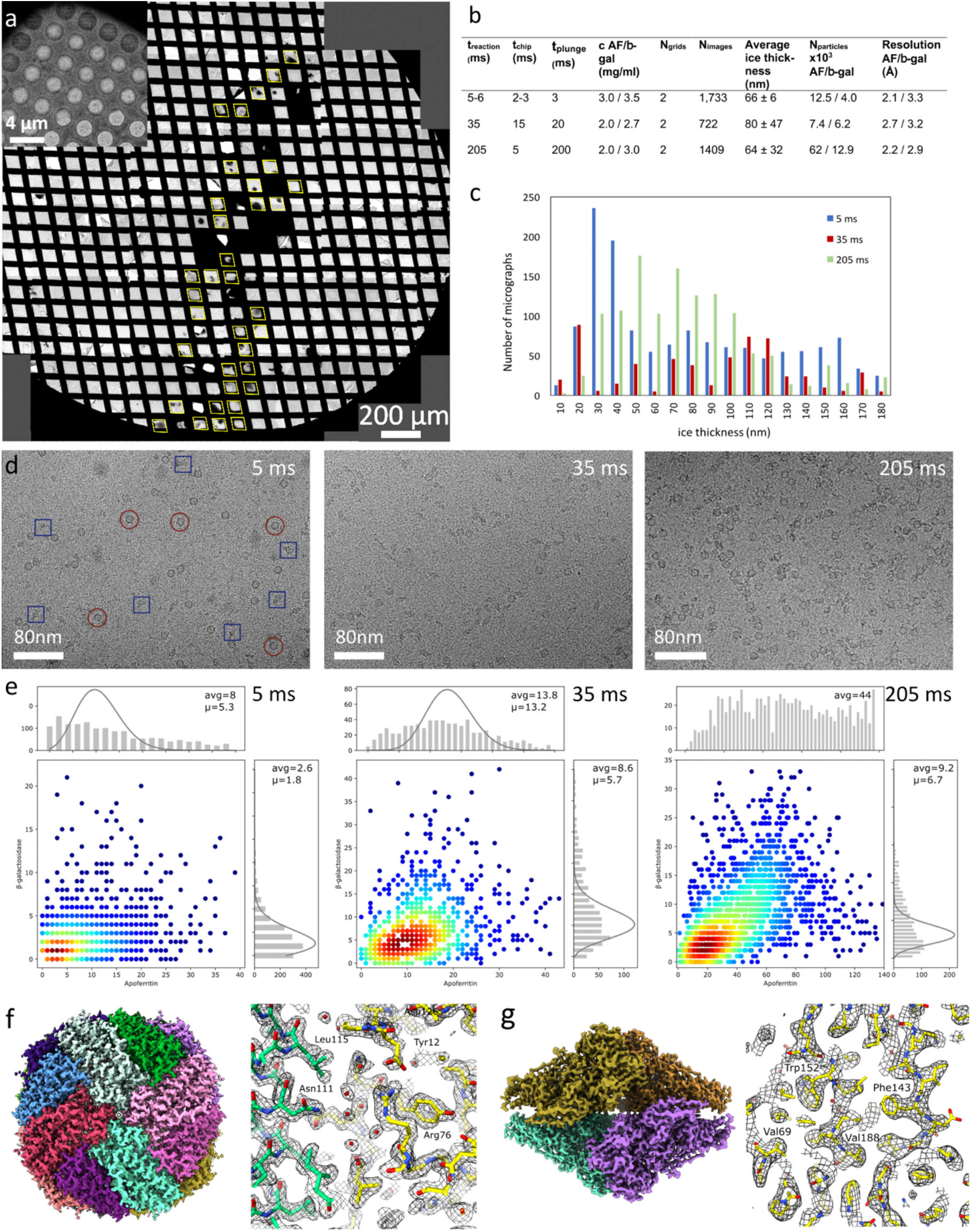
Benchmark of time-resolved cryoplunger. **a**, Atlas of an EM grid with sprayed apoferritin and β-galactosidase sample plunged with a delay time of 200 ms. Grid squares labeled in yellow indicate positions suitable for single particle data collection. The grid was plunged with a laser frequency of 3500 Hz using a chip with 2 nozzles. The inset in the top left is an example of a droplet spread on a holey grid covered with thin continuous carbon support. **b**, Conditions, and statistics of trEM grids and EM reconstructions of apoferritin (AF) and β-galactosidase (β-gal). **c**, Distribution of ice thickness in the holes for the grids with mixed apoferritin and β-galactosidase. **d**, Representative images for “reaction times” of 5, 35, and 200 ms show mixed protein particles with high contrast. In the 5 ms image selected apoferritin and β–galactosidase particles are indicated by red circles and blue squares, respectively. **e**, Distributions of apoferritin and β-galactosidase particles on the micrographs used for high-resolution reconstruction for three time points. Histograms for both types of particles are shown and fitted with the Poisson distribution, except for 200 ms apoferritin, for which no meaningful fit was obtained. **f, g,** 3D maps of apoferritin and β-galactosidase at a resolution of 2.1 and 3.3 Å respectively, reconstructed from a grid plunged with a “reaction time” of 5 ms.

The number of droplets observed on the EM grid atlases was consistent with the expected number of sprayed microjets (**Fig. 2a**), and it increased with increasing plunge time because the grid resides longer in front of the nozzle (**Fig. 1e**). Many droplets deposited on the EM grids spread producing areas of thin ice suitable for high-resolution protein imaging (**Fig. 2**). We observed that the droplets spread fast on the surface of plasma-cleaned carbon holey grids, but many holes remained empty (**Supplementary Fig. 5a**). We assumed that this effect is caused by additional surface energy associated with droplets spreading over holes, and it can be mitigated by using a support film. After testing several support films, we found that freshly plasma-cleaned grids with an additional 2-3 nm thick carbon film improved droplet spreading and increased the number of holes suitable for cryo-EM data collection by a factor of 2 to 3 (inset **Fig. 2a**, **Supplementary Fig. 5b**).

Areas with ice thickness below 100 nm were found on grids plunged with plunge times between 3 and 200 ms (**Fig. 2b, c**), and between a few hundred and a few thousand high-contrast images were collected per grid (**Fig. 2b**).

For each time point, cryo-EM data were collected from two EM grids, and 3D maps of apoferritin and β-galactosidase were reconstructed to resolution between 2.1 and 3.3 Å (**Fig. 2b-g, Supplementary Fig 6, Supplementary Table 1**).

The mixing efficiency was assessed from the EM images by analyzing the distribution of the number of apoferritin and β-galactosidase particles per micrograph (**Fig. 2e**). For all timepoints unimodal distributions consistent with complete mixing were observed. Interestingly, when the distributions of particle number per micrograph were analyzed for each protein independently, they were not following a Poisson distribution (**Fig. 2e)**, expected for ideally behaving particles. Rather the distributions had a flat shape which suggested a more complex behavior.

As plunge times decreased, the particle distributions approached a Poisson distribution, which is particularly evident for β-galactosidase, whereas the average number of particles per micrograph decreased (**Fig. 2e**). The observed behavior had been attributed to protein interaction with air-water interface^43,44^, and our results are consistent with other observations suggesting that faster plunging improves protein behavior in cryo-EM grids^43,45^. In contrast, the distribution of particle orientations, while changed with reduced plunge times, did not become homogeneous even at 3 ms plunge time (**Supplementary Fig. 6**), pointing towards a higher complexity of protein behavior on the EM grids than simple adsorption on the air-water interface ^44,45^.

The experiments with the test proteins demonstrated the feasibility of trEM with 5 ms time resolution and high spatial resolution using the developed instrument. To further validate the method, we applied it to study a dynamic protein system.

### trEM on GroEL-GroES complex

The time-resolved plunger was used to study the structural dynamics of the well-characterized GroEL:GroES chaperonin complex. GroEL is an ATPase. It is a homo 14-mer organized in two stacked heptameric rings that facilitate the folding of substrate proteins assisted by GroES homo heptamer which binds to epical domains of GroEL^46,47^. Fluorescence and FRET kinetics experiments estimated that upon binding of ATP, GroEL undergoes conformational changes within a few milliseconds^48^, followed by a much slower complexation with GroES. The bound ATP is hydrolyzed on a timescale of seconds, after which nucleotide releases and GroES dissociates from GroEL with a characteristic time of around 20 s^48,49^.

The trEM experiments were performed by supplying GroEL (5-6 μM) into one channel of the microfluidic chip and a solution containing GroES (10-12 μM) (GroES was not present at 13 ms time point), ATP (4 mM), and amaranth dye (26-38 mM) into the other. The solutions were mixed with an approximate 1:1 (v/v) ratio and plunge-frozen with reaction times of 13, 50, and 200 ms. A steady state continuous turnover reference dataset was collected from a cryo-EM sample plunged using a conventional plunger at around 20 s after initiation of the reaction (**Table 1**).

**Table 1.** Properties of trCryo-EM grids plunged with GroEL, GroES, and ATP.

| $t_{\text{reaction}}$<br>(ms) | $t_{\text{chip}}$<br>(ms) | $t_{\text{plunge}}$<br>(ms) | $C_{\text{GroEL}}$ (mg/ml) /<br>$C_{\text{GroES}}$ (mg/ml) /<br>$C_{\text{ATP}}$ (mM) | $N_{\text{grids}}$ | $N_{\text{images}}$ | Ice thick-<br>ness (nm) | $N_{\text{particles}}$ | Resolution<br>range (Å) |
| --- | --- | --- | --- | --- | --- | --- | --- | --- |
| 13±3 | 5-10 | 4 | 8 / 0 / 4 ** | 15 | 2654 | 143±72 | 58 k | 3.2-4.4 |
| 50 | 15 | 35 | 8-11 / 2 / 4 | 5 | 1782 | 92±62 | 54 k | 2.7 -7.1 |
| 195±5 | 15-20 | 170-185 | 8-9 / 1.5-2 / 4 | 8 | 6497 | 94±78 | 384 k | 2.3-6.3 |
| 2 10<sup>4</sup> | - | 2 10<sup>4</sup> | 3.7 / 0.7 / 2 * | 2 | 4195 | 82 ±23 | 252 k | 2.3-3.4 |

\* For the 20 s time point the final concentrations are given, whereas for other time points the concentrations before mixing. \*\* The 13 ms time point was prepared without GroES because on this time scale yield of complexation is very low.

Similar to the benchmark, for all time points, the droplets generated by LIC spread on the grids coated with thin continuous carbon layer producing areas suitable for high-contrast cryo-EM imaging (**Table 1**, **Supplementary Fig. 7**). Thousands of micrographs usable for 3D reconstructions were collected per grid for 200 ms reaction times, whereas hundreds from the grids plunged with 50 and 13 ms reaction times. The resulting datasets contained 54 to 384 thousand particles (**Table 1, Supplementary Table 2**).

The differences in the composition of the complexes were already visible in the 2D classes (**Supplementary Fig. 7**). Under steady state conditions most of the side views corresponded to football-shaped GroEL:2GroES complexes, whereas at 200 ms, the side views were dominated by GroEL particles with a smaller fraction of GroEL:GroES and GroEL:2GroES complexes. At 50 ms, GroEL represented the major fraction of the particles, while GroEL:2GroES complexes were absent.

To obtain 3D reconstructions, the particles extracted from all time points were combined, and 3D classified to separate different complexes and conformations present in the dataset. The sets of particles assigned to different complexes and conformational states were then split into subsets, each corresponding to a reaction time point, after which individual 3D reconstructions were refined (**Supplementary Fig. 8**)^11,50^.

Fifteen reconstructions were obtained at a resolution between 2.2 and 7.1 Å (**Figs. 3 and 4, Table 1, Supplementary Table 2**) and included 6 different species. In three of those complexes: GroEL-ATP, GroEL-ATP:GroES, GroEL-ATP:2GroES, the phosphorylation state of bound nucleotide was unambiguously resolved. In the other three complexes: GroEL-(ADP), GroEL-(ATP)/(ADP), and GroEL-ATP/(ADP):GroES, the conformations of the GroEL heptameric rings corresponded to ATP- and ADP-bound states, however, the phosphorylation states of the bound nucleotides were ambiguous due to limited resolution and are indicated by round brackets. For all time points, the data were of high-enough quality to enable not only high-resolution reconstructions but also 3D classification and separation of minor populations containing as few as 4 % of particles (see **Fig. 3,** 50 ms timepoint GroEL-(ADP) reconstruction). Interestingly, because of higher fractions of particles of GroEL-ATP complex at 50 and 200 ms than at 20 s the resolution of the reconstructions obtained from the time-resolved data (2.7 and 2.3 Å) is higher than that obtained from the continuous turnover data (2.8 Å).

**Figure 3.**
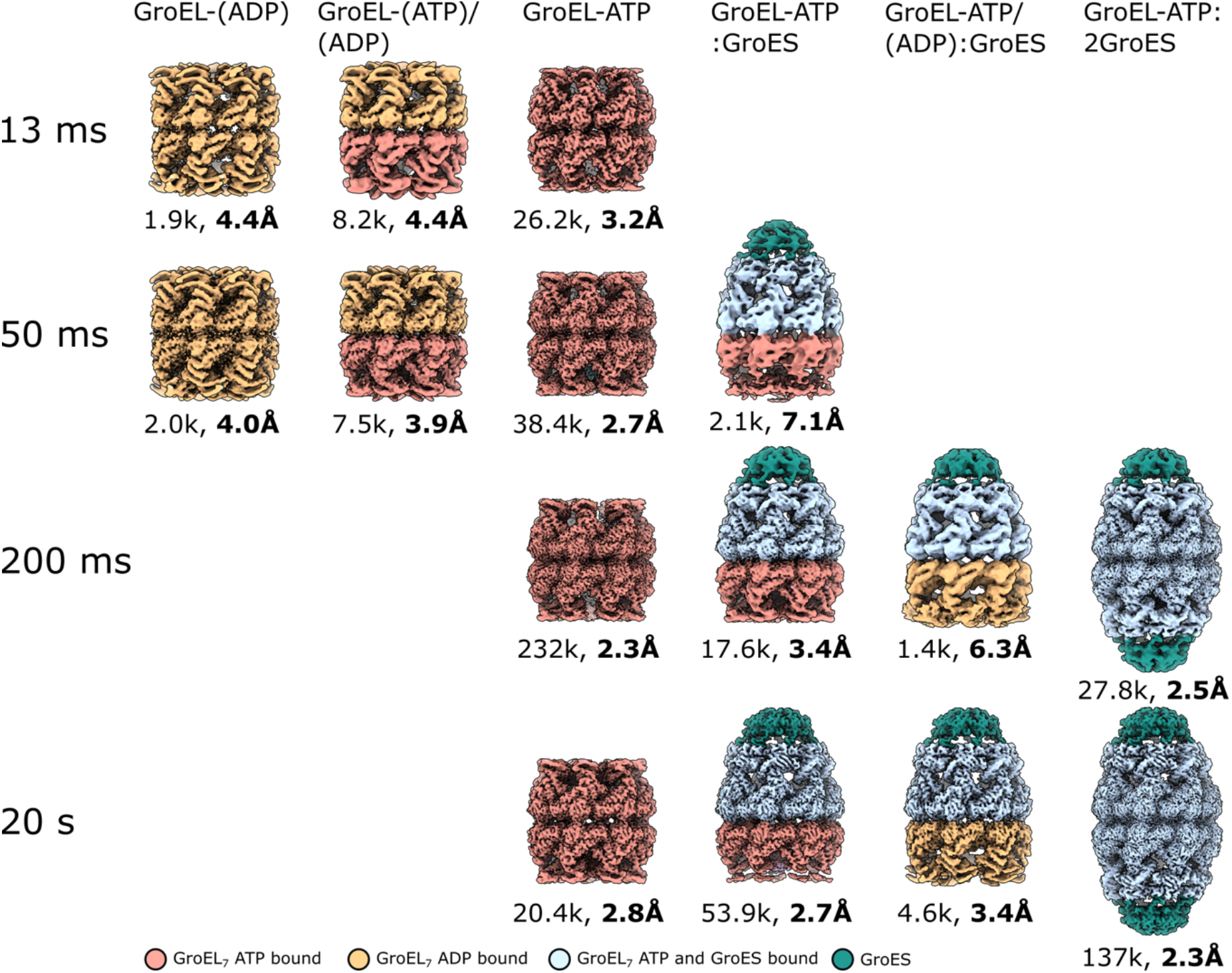
Three-dimensional reconstructions of GroEL:GroES complexes from time-resolved and steady state data. The surface representations of 3D maps reconstructed from trEM data collected from grids plunge frozen with reaction times indicated on the left. The number of particles and gold-standard resolution are indicated under each reconstruction. The density surfaces are colored by protein type and protein conformation: GroES - teal, GroEL in GroES-bound conformation – light blue, GroEL-ATP – brown, GroEL-(ADP) – dark yellow. The nucleotide names in brackets indicate that heptameric GroEL ring conformation corresponds to a certain nucleotide state, but the phosphorylation state, ATP versus ADP, was ambiguous. trEM experiments for 13 ms datapoint were conducted without GroES.

**Figure 4.**
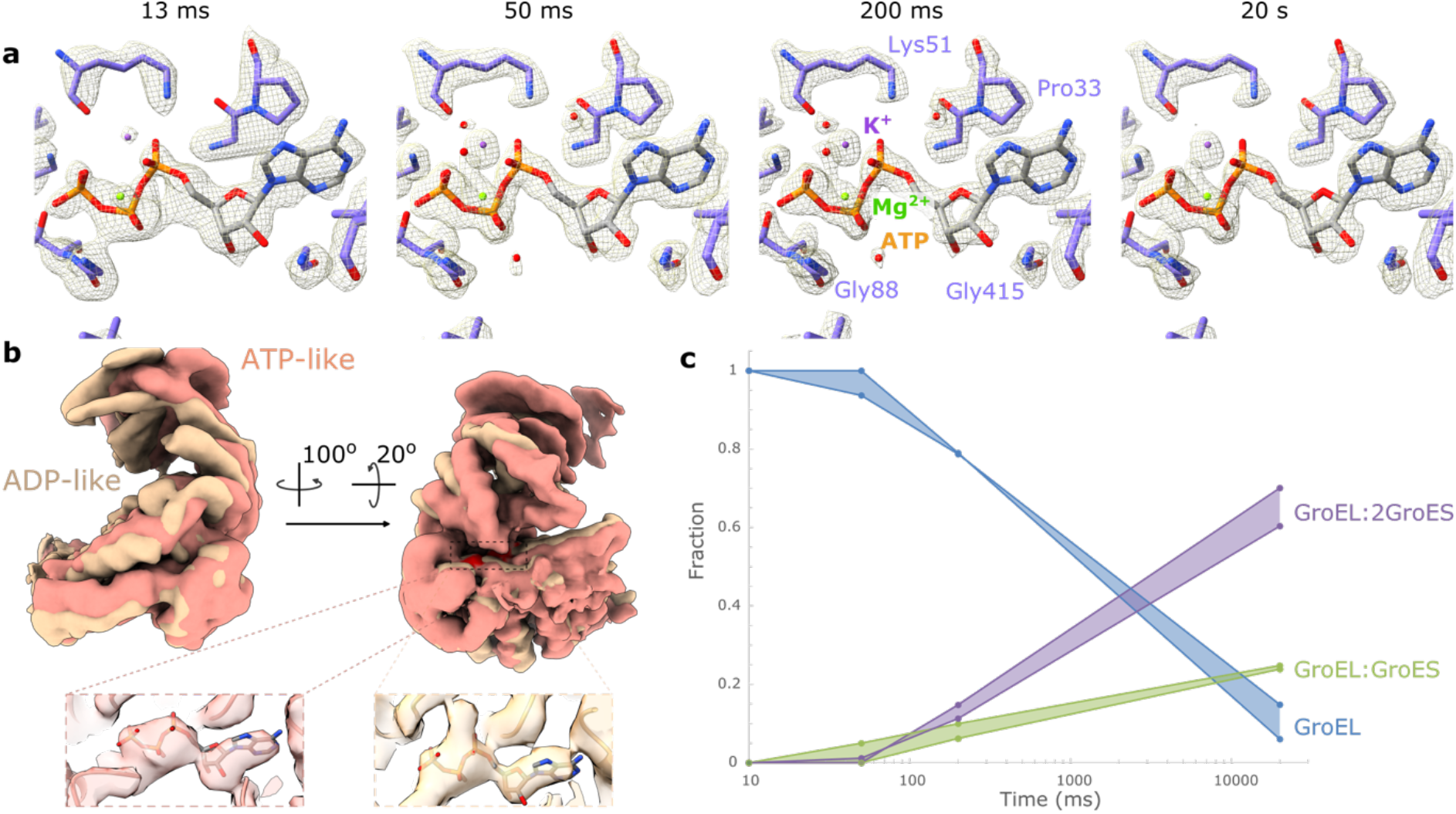
Details of trEM GroEL:GroES reconstructions. **a**, EM maps for GroEL-ATP states are shown around ATP molecule for reconstructions obtained from individual reaction delay times. **b**, Maps of GroEL single subunits in GroEL-(ATP) and GroEL-(ADP) states extracted from the reconstructions of GroEL-(ATP)/(ADP) and GroEL-(ADP) maps calculated from pooled 13 and 50 ms datasets (**Supplementary Fig. 8**). Densities of heptameric rings were aligned then subunits were isolated and low pass filtered for clarity. The red zone shows the surface of bound nucleotide density. Insets show that in both ATP-like and ADP-like rings density of bound nucleotide is visible. The color code of conformations is like in **Fig. 3**. For the ATP-like model, the PDB model with ID 4AAQ was a rigid body fit into the map. For the ADP-like model, the PDB model with ID 4KI8 was used. For representation, ATP molecule (instead of ADP from the PDB 4KI8) was fit into the density for ADP-like conformation **c**, Evolution of fractions of GroEL:GroES complexes determined from trEM data. The fractions of conformations for each time point were calculated by two methods: (1) using a fraction of particles classified into a reconstruction of a specific complex during the 3D classification of pooled data (see Methods for details), (2) using a fraction of particles contributing to high-resolution reconstructions (> 8 Å).

The trEM reconstructions show how the conformational landscape changes as the reaction proceeds (**Fig. 3**). At low reaction times, 13 and 50 ms, apart from the dominant form of GroEL-ATP complexes, GroEL-(ADP) and GroEL-(ATP)/(ADP) complexes are resolved that may represent short-lived states on the way towards the formation of GroEL-ATP conformation. Even though the resolution of these reconstructions will have to be improved to unambiguously resolve the phosphorylation states of the bound nucleotide, they clearly resolved ATP-like and ADP-like conformations of GroEL heptameric rings and nucleotide-binding pocket contains density consistent with a bound nucleotide (**Fig. 4b**).

The density for nucleotide bound to GroEL is unambiguously resolved in all reconstructions, including short time points. More specifically, in GroEL-ATP state at 13 ms time point, the conformation of ATP molecule including adenine ring, three phosphate groups, and coordinating Mg^2+^ and K^+^ ions is well resolved, whereas starting from 50 ms reconstructions obtained at resolutions between 2.3 and 2.8 Å also resolved water molecules in the ATP binding pocket (**Fig. 4a**). These results highlight the utility of the method for investigating the interaction of small molecules with the proteins on a millisecond timescale.

The availability of conformational distributions over the span of reaction time allows us to analyze the kinetics of GroEL:GroES complex formation **(Fig. 4c)**. The fraction of GroEL:GroES and GroEL:2GroES complexes grows between 50 ms (5 and 0 %) and 20 s (~24 and 65 %). The assembly begins with the formation of the GroEL:GroES complex (**Fig 3**, 50 ms data point) followed by the addition of a second GroES heptamer. The major increase in the fraction of the complexes occurs between 200 ms and 20 s which is consistent with the expected rate of GroEL:GroES complex formation estimated by FRET experiments to be in the sub-second range^7^. This demonstrates that cryo-EM samples prepared by our time-resolved plunger qualitatively reproduce complex formation kinetics observed in solution.

The example of GroEL:GroES shows that using our method for preparation of trEM grids, cryo-EM data can be obtained that enable studies of protein-substrate and protein-protein complex formation at near-atomic resolution and resolving structural kinetics with a few milliseconds time-resolution.

## Discussion

This work presents the design, realization, and validation of an instrument for the preparation of time-resolved cryo-EM grids. It is built upon the advantages of a picoliter droplet microfluidic mixer and tunable on-demand generation of microjets by LIC. This approach solves the limitation of other sample preparation methods for time-resolved cryo-EM; it reduces sample consumption and increases the reproducibility of prepared cryo-EM grids while maintaining high time and spatial resolutions.

### Complete mixing with high time-resolution and low sample consumption

The methods of trEM sample preparation can be divided into mixing-based, and flash-photolysis based. The latter relies on the availability of caged compounds or native photosensitivity of proteins and in principle is technically easier to realize. The availability of caged compounds, the complexity of uncaging reaction^51^, and light-induced grid heating^52^ are however among the bottlenecks limiting their widespread applications.

The mixing-based methods, while being technically more demanding, are simultaneously more generic. They allow reactions to be initiated by mixing macromolecules with proteins or small molecules, diluting the sample, and changing salt concentration or pH, for example.

The core advantage of our method stems from combining a droplet-based microfluidic advective mixer with a drop-on-demand spray generator. The chaotic advective droplet-based mixer initiates the reaction in the separated compartments of picoliter drops, where the mixing is defined by the geometry and length of the channel^34^, rather than being dependent on flow instability requiring high flow rates as in chaotic mixers^24,31^. This enables the presented device to operate at low pressure (below 1 bar) and low flow rates reducing sample consumption to around 100 nL per EM grid. When compared to trEM grid preparation by Spotiton^32^ which uses a similar to Berriman & Unwin^21^ approach of on-grid mixing, our method produces a homogeneous mixture of protein with actuating molecules, thus eliminating concentration gradients which can complicate the data analysis.

We demonstrated that our instrument produces reliable and complete mixing of large macromolecules with a residence time down to 2 ms and generates cryo-EM grids with a plunge time down to 3 ms that are useful for high-resolution 3D reconstructions (**Fig. 2**).

The limit sample consumption in the described instrument is in order of a few nanoliters per grid. It is defined by the chip volume, residence time, and time of jetting on the grid. In practice, the consumption is somewhat higher due to the lag time needed to initiate flow in the chip, but still remains below 100 nL per grid, and tens of EM grids are routinely plunged using a volume of 20 μL of protein solution. This includes dead volume in the tubings and protein consumed during optics alignment procedures.

### Controlled preparation of cryo-EM grids

Reproducible spreading of micro droplets into a liquid film of thickness below 100 nm within the millisecond timeframe is the challenge in the mixer-based trEM methods^23,31,53,54^.

Spraying microdroplets generated by a gas-assisted atomizer was among the first approaches to successfully vitrify water for cryo-EM^53^ and was later applied to trEM^21,23–25,31^. Smaller droplets (diameter < 10-20 μm) spread rapidly on hydrophilic surfaces^23,54^, whereas larger droplets, like those generated by Spotiton or Shakeitoff devices^55,56^, required wicking to obtain thin ice. The LIC has the advantage of generating reproducible microjets that disintegrate into smaller droplets, which even at 3 ms plunge time spread into thin layers allowing reconstructions of apoferritin to 2.1 Å resolution (**Supplementary Video 1, Fig. 2**). Depending on the plunge time, between 6 and 70 jetting events occur during the passage of a grid in front of the nozzle. The resulting cryo-EM grids contained between a few tens and a few thousands of holes suitable for high-resolution imaging. The surface usable for data collection can be further increased by improving control over the spreading of the microdroplets and by increasing the frequency of the laser, limited to 5 kHz in the present device.

### Recovering reaction kinetics and conformational landscape

Kinetics of conformational transitions constitutes the basis for understanding protein function. Time evolution of protein conformations and composition can be used to derive the kinetic parameters from the trEM experiments^11,20^. The systematic trEM experiments with a few milliseconds time resolution that enable kinetics analysis remain challenging and were reported only for ribosomes^20^. We show that our instrument can deliver high-resolution cryo-EM data down to 5 ms reaction time enabling up to 2.1 Å resolution for apoferritin and 3.3 Å for β-galactosidase and GroEL. It captures protein complex formation kinetics consistent with FRET data. trEM grids with delay times covering the span of a reaction can be prepared in one session by adjusting the flow rates and plunge times, thereby increasing the consistency of kinetics data and reducing sample consumption.

It is worth noting that in trEM reaction proceeds under two different conditions: in the microchannel of the microfluidics chip and on the grid within a hundred nanometer thin film, which may influence the kinetics of reactions and will require further experimental studies.

### Towards microsecond time resolution

The physics-imposed limits of time resolution in trEM are defined by the sample cooling time estimated to be in the order of 100 μs^15^ or faster^57^. In our plunger, the resolution is limited by the chip residence time and plunge time. The limits are technical and can be reduced below 1 ms. Mixing times below 500 μs have already been demonstrated for droplet-based mixers^34^, whereas the plunge time is limited by the frequency of jetting rather than the velocity of the plunger arm. We envision that further technical improvements to the setup will achieve a time resolution of below 1 ms.

The described method for preparation of time-solved cryo-EM grids, combined with more intelligent and automated cryo-EM data collection schemes, and methods enabling more automated processing of strongly heterogeneous single particle cryo-EM data, will enable routine structural studies of processes taking place in proteins on a millisecond time scale. This will further our understanding of the function-related conformational dynamics of proteins and biological molecular machines in particular.

## Methods

### Microfluidic chip fabrication

The microfluidic chip has been fabricated using a standard soft lithography technique^58,59^. The polymeric part of the chip was fabricated as a replica of a mold obtained by photolithography. The designed pattern was printed on a transparency mask with a resolution of 6 μm (Selba SA) and transferred on a silicon wafer coated with a photoresist. The photoresist (SU8 3050, Kayaku Advanced Materials, Inc.) was spin-coated at 3000 rpm for 30 s, using a Polos spin-coater (SPIN150i, SPS-Europe B.V.) to achieve a thickness of 50 ± 5 μm, followed by a soft-bake step on a hot plate for 15 min at 95 °C. Then the pattern was transferred by illuminating the photoresist through the photomask with UV light at 365 nm and power of 30 W for 7 s using mask aligner UV-Kube 3 (Kloé). The post-bake step was performed by placing the wafer on the pre-heated hot plate at 95 °C for 10 minutes. Then the un-exposed areas were dissolved with SU8 developer solution (Kayaku Advanced Materials, Inc.) for 8 minutes. During this step, the developer solution was continuously gently mixed on a shaker to ensure the removal of unexposed material from the pillar area. The resulting master was examined by an optical profilometer (Profilm3D, Filmetrics) to verify the structure details and the resist thickness. Next, the microchannels were formed in polydimethylsiloxane (PDMS, Sylgard 184 Dow Corning) through the casting of the uncured polymer against the master. The polymer base was mixed with a curing agent in a 10:1 ratio, to activate the cross-linking process. The mixture was poured on the master mold and then degassed in a desiccator for about 30 minutes or until all the air bubbles were removed. The curing was finalized by placing the polymer mold in an oven at 65 °C overnight. Then, the PDMS was detached from the master mold and access holes for the tubing connections were perforated using a surgical puncher with a diameter of 1.5 mm (Rapid-Core sampling tool, Electron Microscopy Sciences). In addition, the PDMS chip was cut at approximately 200-400 μm from the end of the edge of the droplet merging region transversal to the outlet channel to make the nozzle openings. Finally, the polymer device was sealed with a glass slide after exposing both PDMS and glass surfaces to oxygen plasma in plasma cleaner Cute (Femto Science) at base pressure 0.8 torr, power 100 W, process pressure 0.5 torr, and duration 1 minute. The glass slide with a thickness of 170-190 μm (Ted Pella, Inc.) was carefully manually aligned to the PDMS part of the chip under a stereo microscope to obtain a sharp rectangular opening in the chip. An average residual offset between aligned surfaces was estimated to be below 5 μm. Using thin glass facilitated the laser alignment and focusing during the chip operation. The assembled chip was kept on a hot plate at 180 °C for a minimum of 4 hours to render the PDMS surface of the channels hydrophobic, which was required for the stable formation of water in oil droplets.

### Time-resolved plunger setup

The plunger device was assembled around a CERNA microscope (Thorlabs, Inc.) with a 5x objective (LMH-5X-532, Thorlabs Inc., **Supplementary Fig. 1a, b**). A high-speed camera Phantom VEO410L was mounted on the upper port of the trinocular (WFA4001, Thorlabs, Inc.). A custom-designed dewar with thermostated ethane vial controlled by CN63200-R1-LV controller (Omega Engineering, Inc.) was placed under the objective on a manual XY stage (XYT1, Thorlabs Inc.). The microfluidic chip was placed on a 3D printed holder mounted on a manual Z and motorized XY stage (MVS1, PLS-XY, Thorlabs, Inc). The grid plunger was designed to operate in the confined space of the microscope. An aluminum arm with a steel clip (Micro Serrefine Clamp, 18055-04, Agntho’s AB), used to mount EM grid, affixed to its end, was actuated by a rotational voice coil (extracted from Hitachi Deskstar HDS721010KLA330 hard drive). An incremental encoder 500 ppR (AEDB-9140-A13, Broadcom, Ltd.) was mounted coaxially with the arm (**Fig. 1d**). A plunger arm driver capable of supplying a pulse width modulated (PMW) signal with voltage amplitude between 5 and 220 V to the voice coil was designed. It enabled accelerating the arm in both directions of rotation and was coupled to a microcontroller (Arduino MEGA-2560) that was used to accelerate the arm in a position-dependent manner. A laser beam from the second harmonic of a Q-switched Nd:YAG laser (AO-L-532, CNI Optoelectronics Tech. Co., Ltd.) with an emission wavelength of 532 nm and pulse duration of 6 ns was expanded using a 10x beam expander (BE10-532 10X, Thorlabs, Inc.). The laser beam was then directed with a system of mirrors (**Supplementary Fig. 1a, b**) towards the side entry port of the microscope and brought onto the optical axis of the objective lens by a beam splitter (BSW4R-532, Thorlabs, Inc). A diffractive beam splitter (MS-214-Q-Y-A, HOLO OR, Ltd.) introduced to the laser path (**Supplementary Fig. 1a**) was used to split the laser beam into six equiangularly spaced beams. The exposure of the microfluidic chip to the laser beam was controlled by a shutter (SH05R, Thorlabs, Inc.) coupled with the Arduino board. The laser beam intensity was continuously monitored by a thermal power meter (S401C, Thorlabs, Inc.). The plunger operation was controlled using a program written in LabView 2020 (National Instruments).

### Operation of the time-resolved cryo-EM plunger

The microfluidic chip was operated by applying positive pressure to the fluidic channels using a pressure controller (LineUP Flow EZ, Fluigent) connected to each of the chip inlet channels. The flow in the microfluidic chip was started by first setting the pressure values for each channel and then simultaneously initiating the flow. An additional pressure controller (LineUp Push-Pull, Fluigent), applying negative pressure was connected to the oil extractor channels. Depending on the required residence time, the applied pressures varied in a range of 60 to 400 mbar for the aqueous and oil solutions, and a range of −10 to −100 mbar for the extractor channels. During the experiments, the Q-switched laser was operated at a frequency in the range of 3500-5000 Hz and energy per pulse between 50 and 120 μJ. The resulting laser energy at the surface of the chip per focal spot was 4-10 μJ/pulse. To enable laser-induced cavitation, Amaranth dye (CAS Nr 915-67-3, Sigma-Aldrich) was added at a concentration of 20 mg/mL (36 mM) to the solution fed into one of the aqueous channels. The water in oil droplets were formed using a mix of fluorinated oil (3M, Novec 7500 Engineered Fluid) with 5 % fluoro-surfactant 1H,1H,2H,2H-Perfluoro-1-octanol (CAS Nr 647-42-7, Sigma-Aldrich).

The plunge times were measured for each grid plunging using an Arduino timer which registered times for the rotation positions that corresponded to the grid in front of the nozzle and at the ethane surface. The residence time was measured from high-speed videos by visually following aqueous droplets from the droplet formation until the ejection.

The trajectories of the plunger arm were analyzed using PCC v3.7 software (Phanthom, USA).

### Production and purification of proteins

#### Apoferritin

Apoferritin was produced and purified as described elsewhere^60^.

#### Beta-Galactosidase

After isolation of total bacterial DNA, a construct was cloned by PCR from the LacZ gene of *Escherichia coli* MG1655 with a His-tag on the C terminus by Gibson Assembly (New England Biolabs) in a pET-28b vector using the following primers LacZ200 (TAACTTTAAG AAGGAGATAT ACCATGACCA TGATTACGGA TTCACTGGCC GTCGTTTTAC) and LacZ201 (GGTGCTCGAG TGCGGCCGCA AGCTTATTAT TAATGATGAT GATGATGATG GCCTTTTTGA CACCAGACCA ACTGGTAATG).

After the transformation of the plasmid in BL21 *E. coli,* cells were grown in Luria–Bertani medium at 37 °C, with 50 μg ml^-1^ kanamycin, to an OD at 600 nm of 0.7. The protein expression was induced with 1 *mM* IPTG (Inalco) for 4 h. Subsequent steps were performed at 4 °C. The cells were harvested by centrifugation at 6500 *g* for 15 min and resuspended in 25 ml lysis buffer per pellet of a liter of culture (50 mM HEPES–NaOH pH 8, 200 mM NaCl, 1 mM MgCl_2_), supplemented with 50 μg ml^-1^ DNase I (Merck/Sigma–Aldrich) and 1 mg ml^-1^ lysozyme (Merck/Sigma–Aldrich). After 1 h incubation while stirring, cell debris was spun down at 39 000 *g* for 30 min. The supernatant was loaded on a 5 ml HisTrap HP column (Cytiva) equilibrated with equilibration buffer (25 mM HEPES pH 8, 100 mM NaCl, 20 mM Imidazole, 1 m*M* tris(2-carboxyethyl) phosphine (TCEP)). The protein was eluted from the column with an equilibration buffer supplemented with 500 mM imidazole. Overnight dialysis against the SEC buffer (25 mM HEPES pH 8, 100 mM NaCl1 mM TCEP) was used to remove the remaining imidazole. The dialyzed sample was concentrated to approximately 4 ml, with a 30 kDa cut-off Amicon Ultra Centrifugal Filters (Merck) and ran on a size-exclusion Superdex 200 increase 10/300 GL column (Cytiva) equilibrated in the SEC buffer. The peak fractions containing pure beta-galactosidase were concentrated and stored at −80°C after flash freezing in liquid nitrogen.

#### GroEL and GroES

A synthetic construct was made, based on the GroEL gene of *E. coli* MG1655, in a pET-15b vector (Genscript). After the transformation of the plasmid in BL21 *E. coli,* cells were grown in Luria–Bertani medium at 37 °C, with 100 μg ml^-1^ carbenicillin, to an OD at 600 nm of 0.4. The protein expression was induced with 0.5 mM IPTG (Inalco) for 5 h. The supernatant was obtained as described above with a slight difference in buffer composition (50 mM Tris-HCl pH 8, 1 mM EDTA, 20 mM MgCl_2_, 1 mM TCEP). After centrifugation, ammonium sulfate was added to the supernatant to a final concentration of 40 % (*w*/*v*). This suspension was centrifuged at 39 000 *g* for 1 h and supernatant was collected. Next, the concentration of ammonium sulfate was raised to 50 % (*w*/*v*) and another round of centrifugation was performed. The pellet was resuspended in 6 ml purification buffer (50 mM Tris-HCl pH 8, 1 mM EDTA, 1 mM TCEP). Ammonium sulfate was removed by overnight dialysis against the purification buffer. The dialyzed sample was diluted with purification buffer to a conductivity of 10 μS/cm and loaded on a 24 ml Source Q15 Anion Exchange (Cytiva) column pre-equilibrated with the purification buffer. The protein was eluted by a linear gradient of the purification buffer supplemented with 0.6 mM NaCl from 0 to 100 % over 20 column volumes. The peak fractions containing pure GroEL were stored at −80°C after flash freezing in liquid nitrogen.

Before preparation of cryo-EM grids, the thawed GroEL was further purified using Superose 300/10 increase column in 25 mM Tris-base pH 7.5, 0.5 mM TCEP, 50 mM KCl, 10 mM MgCl_2_.

A synthetic construct was made, based on the GroES gene of *Escherichia coli* MG1655, in a pET-15b vector (Genscript). The transformation and protein expression are the same as described for GroEL apart from 1 mM IPTG (Inalco) used to induce protein expression. The supernatant was obtained as described above. The supernatant was loaded on a 24 ml Source Q15 Anion Exchange (Cytiva) pre-equilibrated with the purification buffer (50 mM Tris-HCl pH 8, 1 mM EDTA, 1 mM TCEP). The protein was eluted by gradient elution with the purification buffer supplemented with 0.6 mM NaCl. The peak fractions containing pure GroEs were stored at −80 °C after flash freezing in liquid nitrogen.

Before preparation of cryo-EM grids, the thawed GroES was further purified using Superdex 200/10 column in 25 mM Tris-base pH 7.5, 0.5 mM TCEP, 50 mM KCl, 10 mM MgCl_2_.

### Preparation of cryo-EM grids

For the preparation of cryo-EM grids using the time-resolved plunger, Quantifoil R2/1, holey, 300 mesh grids were coated with a continuous carbon layer of 3 nm in sputter coater EM ACE600 (Leica). The continuous carbon layer was first deposited on mica support of 25 x 25 mm (Cat Nr 71850-11, Electron Microscopy Science). The carbon layer was floated on the water surface and then transferred to the cryo-EM grids by raising them to the water surface under the carbon layer. Before plunging, the grids were glow discharged in vacuum plasma system Cute (Femto Science) for 1 minute at a frequency of 50 kHz, process pressure 0.8 torr, and power 25 W.

Apoferritin in 20 mM HEPES, 100 mM NaCl, pH 7.5, and β-galactosidase in 25 mM HEPES, 100 mM NaCl, 1 mM TCEP, pH 8 were supplied into separate channels. The flow parameters were set to achieve a 1:1 mixing ratio. For other details see **Supplementary Table 3**.

GroEL and GroES were freshly gel filtered as described above. GroEL (8-11 mg/ml) was supplied in one and GroES (1-2 mg/ml), ATP (4 mM), amaranth (26-40 mM), and CHAPS (0.2 %) in another channel of the microfluidic chip. Concentrations of proteins were adjusted to obtain GroEL:GroES molar ratio of 1:2. CHAPS was added to reduce the preferred orientation of GroEL.

For the 13 ms reaction time point, GroEL (8-9 mg/ml) was supplied into one and ATP (4 mM), amaranth (40 mM), and CHAPS (0.2 %) into another channel.

For 20 s reaction time point, GroEL (3.7 mg/ml) was mixed with GroES (0.7 mg/ml), ATP (2 mM), and CHAPS (0.1 %). 4 μl sample was applied on a freshly glow discharged Quantifoil R2/1 copper 300 grid in CP3 (Gatan) under a relative humidity of 90 % and temperature of 22 °C then blotted with Whatman 1 filter paper from both sides for 2.5 s and plunge-frozen in liquid ethane at −181 °C.

### Cryo-EM data collection

Cryo-EM images were collected on a JEOL CryoARM 300 electron microscope operated at 300 kV equipped with a cold field emission gun (cFEG), an in-column Ω energy filter, and a four-lens condenser system^61^. Images were recorded on a K3 detector (Gatan) operating in correlated-double sampling (CDS) mode. Movies consisting of 59 or 60 frames were acquired with a nominal magnification of 60,000, corresponding to a pixel size of 0.73-50.750 Å. The microscope illumination conditions were set to spot size 6 and alpha 1. The energy filter slit width was 20 eV. The exposure time was 2.796 or 3.036 s, corresponding to an average dose per frame of 1 or 1.06 e^-^ / Å^-2^. The defocus varied between −0.5 and −3.5 μm. Automated data collection was performed using SerialEM v 3.8.0. Coma-free alignment was performed before launching the data collection. Five images were collected per single stage position that was set at the center of a grid hole. The ice thickness was measured using the energy filter method as described elsewhere^61^.

### Image processing

#### Image processing of apoferritin and β-galactosidase data

For all three datasets, the collected images were processed as follows. First, the movies were aligned in MotionCor2^62^, and the Contrast Transfer Function (CTF) parameters were determined in CTFFIND 4.1^63^. A subset of 25-30 selected images was used for manual particle picking and training crYOLO 1.7.6^64^. For the three datasets, the total number of picked particles was: 134,934 from 1733 micrographs, for 5 ms; 93,847 from 722 micrographs, for 35 ms; and 260,531, from 1409 micrographs, for 205 ms. The particles were then extracted and processed in Relion 3.1.2^65^, initially in a box size of 90 pixels with four times binned pixels. Apoferritin and β-galactosidase particles were separated after the first round of 2D classification. For the 5, 35, and 205 ms datasets a total of 19,466, 17,348; 23,974, 32,884; and 62,119, 32,414 particles were selected for apoferritin and β-galactosidase, respectively. These subsets were further reduced by selecting particles from good classes after successive 2D classification steps. For both proteins, initial unmasked 3D refinement was performed using an initial model generated from PDB files (PDB IDs: 7A4M, 4V40) and filtered to 20 Å resolution. After the first 3D auto refinement, 3D classification was performed and for the 5, 35, and 205 ms dataset, 12,593, 4013; 7432, 6204; and 60,146, 12,887 particles were retained for apoferritin and β-galactosidase, respectively. Next CTF refinement and Bayesian polishing were performed followed by 3D refinement and postprocessing. Particles were re-extracted in a bigger box size with the size chosen according to the resolution of the refined map and new refinement performed. For the 5, 35, and 205 ms datasets, the final resolutions of 2.1, 3.3 Å; 2.7, 3.2 Å; and 2.2, 2.9 Å for apoferritin and β-galactosidase were obtained, respectively (**Supplementary Table 1**).

#### Image processing of GroEL data

Initial steps of image processing for GroEL:GroES dataset were identical to those described above for apoferritin and β-galactosidase datasets. The particles were auto-picked in Cryolo1.8 and extracted with pixels binned 2 or 4 times. Then, each dataset was processed either individually or combined with data from the same time point. The processing was performed in Relion 3.1 and Relion 4.0-beta^66^. For the 2D classification, mask diameters of 330 and 250 Å were used for the side and the top views of GroEL, respectively. The initial model was generated using a crystal structure of ATP-bound GroEL (PDB ID: 4AAQ^46^) converted to a 3D map and low pass filtered to between 30 and 60 Å.

First, the particles from time points of 50, 200 ms, and 20 s selected after 2D classification were merged, totaling around 700 k particles, and 3D auto-refined with C7 symmetry into a consensus map to 2.6 Å resolution. Next, CTF refinement was performed for symmetric and asymmetric aberrations, anisotropic magnification, and per particle defocus. A subsequent round of 3D auto-refinement in C7 symmetry using a mask generated by filtering the consensus map to 15 Å and extending it by 8 pixels with 8 pixels soft edge, generated a 2.4 Å resolution consensus map. The particles were 3D classified with the mask without aligning the particles, into 20 classes with C1 symmetry. T value of four was used for the classification. Classification generated six good classes which showed secondary structural features of GroEL, corresponding to different states of GroEL and GroEL-GroES complexes (**Supplementary Fig. 8**). Particles from each class were selected and split based on a time point. This generated a total of 18 particle sets. The particle sets were 3D auto-refined individually, resulting in 10 reconstructions at a resolution better than 8 Å.(**Fig. 3, Supplementary Table 2, Supplementary Fig. 7**). The individual subsets obtained from class 3 refined to a resolution below 10 Å and thus were not considered further. Class 18 contained nearly 120 thousand particles in conformation assigned to GroEL-(ATP), but the particle subsets corresponding to individual time points refined to resolution significantly lower than those corresponding to the same state from class 9. Therefore, these reconstructions were not used further. We noticed, however, that the reconstruction of class 18 displayed small differences in the conformations of apical domains when compared to class 9.

Two good maps were obtained from a 50 ms timepoint. The first map, from class 9 assigned to the GroEL-ATP complex, was generated with 38,440 particles and reconstructed to 2.7 Å resolution with D7 symmetry. 2170 particles from class 15 were refined with C7 symmetry and produced a map at 7.1 Å resolution.

Refinement of classes corresponding to the 200 ms datapoint resulted in four reconstructions of different GroEL and GroEL:GroES complexes (**Supplementary Table 2b, Supplementary Figs 7, 8b-d**). Class 9 corresponding to the GroEL-ATP complex, with 231,968 particles, was refined to a resolution of 2.3 Å with D7 symmetry. The other three classes showed one or two GroES bound to GroEL. Classes 4 and 15, comprising 1433 and 17,592 particles were reconstructed with C7 symmetry and resolved to 6.3 and 3.4 Å, respectively. The fourth map generated with 27,784 particles from class 14 represented the GroEL-ATP:2GroES complex, it was refined with D7 symmetry and resolved to 2.5 Å.

Four states of GroEL and GroEL:GroES were refined from data collected at 20 s time point (**Supplementary Table 2b, Supplementary Figs 7, 8b-e**). Particles from classes 9 and 14 were refined with D7 symmetry, whereas particles from classes 4 and 15 were refined with C7 symmetry. Class 9 contained 20,392 particles and resolved to 2.8 Å. Classes 4 and 15 comprising 4612 and 53,885 particles reached resolutions of 3.4 Å and 2.7 Å, respectively. The last map of the GroEL-ATP:2GroES complex, from class 14, was generated from 136,955 particles and refined to 2.3 Å resolution.

Data from the 13 ms time point were initially processed separately from the other time points. 57,914 particles were selected after 2D classification and refined to a consensus 3D map with D7 symmetry, after which the particle set was refined for asymmetric and symmetric aberrations, anisotropic magnification and per particle defocus. 3D auto refinement produced a map at a resolution of 3.8 Å. A mask was generated by filtering this map to 15 Å, extending edges by 8 pixels with soft edges of 5 pixels. The particles were 3D classified in D7 symmetry into 6 classes without alignment with a mask using the T parameter of four. The classification resulted in three classes populated with particles. The first class contained 26,156 particles (45 %) and refined to 3.2 Å. It corresponded to GroEL-ATP conformation, the second class contained 20,675 (35 %) particles and did not refine to high resolution. The third class contained 6480 (11%) particles and refined to 4.2 Å resolution (**Supplementary Fig. 8f**), however, reconstruction from these particles corresponded to an intermediated conformation between GroEL-ATP and GroEL-ADP.

To check if this conformation exists in other datasets, the particle sets from other time points assigned to conformations without bound GroES, classes 9 and 18, were 3D classified separately (for each class and each time point) into 10 classes each in D7 symmetry without aligning particles. All good subclasses originating from class 9 corresponded to the GroEL-ATP conformation. Classification of 50 ms subset from class 18 produced classes corresponding to GroEL-ATP, GroEL-(ADP), and the intermediate conformation.

Classification of 200 ms subset from class 18, containing 84,629 particles, produced classes corresponding to GroEL-ATP and the intermediate conformation. The intermediate conformation comprised 6304 particles which auto-refined to 4.8 Å resolution. Classification of the 20 s subset from class 18 produced only one good class corresponding to GroEL-ATP conformation. The subset of 6304 particles was merged with all particles from 13 and 50 ms time points and classified with C7 symmetry without aligning the particles with the mask in eight classes using the T parameter of four. The classification resulted in four good classes showing the secondary structure of GroEL. In three of them, both heptameric rings were in ATP-bound conformation of GroEL. In one of the classes, both rings were in GroEL-(ADP) conformation. These particles were split per time point and refined in C7 symmetry. This resulted in the GroEL-(ADP) reconstruction at 4.4 Å for 13 ms time point and GroEL-(ATP)/(ADP) conformation at a resolution of 4.4 and 4.6 Å for 50 and 200 ms subsets, respectively. Because of the inconsistency between initial and refined conformations, an alternative classification was performed.

Only particles from 13 and 50 ms time points were pooled together and a consensus map at a resolution of 3.4 Å was obtained with C7 symmetry. A mask was generated by filtering the consensus map to 15 Å and extending edges with soft edges of 8 pixels each was used for 3D classification without alignment into 8 classes and with a T parameter of 4. Three classes with well-defined GroEL features were obtained (**Supplementary Fig. 8g**). Class 2 map corresponded to the symmetric GroEL-ATP conformation. The class 1 map had one heptameric ring in GroEL-(ATP) and another in GroEL-(ADP) conformation [GroEL-(ATP)/(ADP)], whereas reconstruction of class 8 corresponded to symmetric GroEL-(ADP) conformation. The particle sets from GroEL-(ATP)/(ADP) and GroEL-(ADP) classes were split into their respective time points and auto-refined separately. The GroEL-(ATP)/(ADP) subset of the 13 ms time point comprised 8252 particles that refined with C7 to 4.4 Å resolution, and the 50 ms time point comprised 7566 particles that refined with C7 to 3.9 Å resolution. The subset corresponding to GroEL-(ADP) conformations comprised 1947 and 2030 particles in 13 ms and 50 ms time points. The auto refinement in D7 symmetry yielded maps at a resolution of 4.4 and 4.0 Å, respectively. The final maps were sharpened by the Relion post-processing procedure and filtered by local resolution.

The identity of the complex conformations was established using high-resolution maps in which both GroEL conformation and bound ligands were unambiguously resolved. The GroEL-ATP conformation was further confirmed by the rigid body fitting of the GroEL-ATP model with PDB ID 4AAQ. The assignment of GroEL-ADP conformation was based on a comparison of the maps with the structure of GroEL-ADP complex, PDB ID 4KI8.

#### Kinetics of complex formation

The fractions of GroEL, GroEL:GroES, and GroEL:2GroEL complexes at each time point were estimated as follows. After the 3D classification step of combined 50, 200 ms, and 20 s datasets, as described above, of 6 good well-resolved classes, 2 classes corresponded to GroEL molecule, 3 classes to GroEL:GroES complex, and one to GroEL:2GroES. The particles corresponding to each complex were merged and side views with tilt angles between 35 and 135 degrees were retained thus excluding top views poorly discriminating the complexes. Then particles corresponding to each time point and each type of GroEL:GroEL complex were counted and normalized by the total number of particles from the time point. Similarly, after all the classifications, and high-resolution refinements (**Supplementary Figs. 7, 8**), particle fractions of GroEL, GroEL:GroES, and GroEL:2GroES complexes were estimated taking into account only the particles contributing to high-resolution reconstructions (>8 Å, **Supplementary Table 2b**). Particles with tilt angles of 35 to 135 were considered for the calculations.

#### B-galactosidase activity assay

Purified β-galactosidase (2.2mg/ml) was mixed with 40 mg/ml dye prepared in 100 mM NaH2PO4, 10 mM KCl, and 1 mM MgSO4 in 1:1 volume proportion such that the final concentration of protein was around 1.1 mg/ml. The mixture was passed through the microfluidics chip under conditions identical to those used for time-resolved plunging of the cryo-EM grids, i.e., the protein solution was encapsulated in the droplets, the droplets were merged, and the spray was generated using laser-induced-cavitation. The experiments were performed at different residence times in the microfluidics chip. The energy of the laser was set to 50 μJ/pulse and the frequency varied between 1500 and 5000 Hz. The ejected spray was collected for a range of protein retention times. To ensure that the proteins were exposed to only one laser pulse, pressure on the channels carrying the fluids and the frequency of the laser were adjusted such that the fluid shifted significantly between the sequential laser pulses maximizing the ejected volume per laser pulse. Collected samples were diluted 20 times with reaction buffer A (100 mM NaH_2_PO_4_, 10 mM KCl, 1 mM MgSO_4_).

The activity of β-galactosidase was measured using a spectrophotometric assay in which o-Nitrophenyl-β-D-Galactopyranoside (ONPG, CAS Nr: 369-07-3 Calbiochem Sigma-Aldrich) was used as a substrate. B-galactosidase converts ONPG into galactose and *o*-nitrophenol, out of which, *o*-nitrophenol absorbs at 420 nm. The concentration of the product was followed on UV-Visible spectrophotometer Cary 100Bio (Varian) at the wavelength of 420 nm for 5 mins in polystyrene cuvettes with 1 cm optical path length at 25° C. The reaction buffer included 855 μL of buffer A to which 45 μL of 1 M β-mercaptoethanol, and 20 μLof 4 mg/ml of ONPG were added. The reaction was initiated by the addition of 10 μLof protein sample. The reaction was repeated thrice for each collected sample of corresponding mixing time. The rate of reaction was estimated from the linear interpolation of the absorption curve in the time interval between 0.15 and 0.6 min.

#### Optical measurement of mixing in microfluidics channels

The mixing efficiency inside the aqueous droplets in oil in the microchannel was optically assessed by recording fast movies of the chip operating at varied fluid flows. The experiment was performed using water, a 16 mM Direct red 81 (CAS Nr 2610-11-9, Sigma Aldrich) or Amaranth dye solution (CAS Nr 915-67-3, Sigma Aldrich), and fluorinated oil as continuous phase (3M, Novec 7500 Engineered Fluid), to which a 5 % of fluoro-surfactant was added (1H,1H,2H,2H-Perfluoro-1-octanol, CAS Nr 647-42-7, Sigma-Aldrich).

The mixing efficiency was estimated after each bending region of the channel, by evaluating the distribution of the pixel intensities inside the droplets at the sequential turns of the serpentine. Movies were recorded using a 5x objective (Thorlabs LMH-5X-532) and a highspeed camera (Phantom VEO MODEL 410L). A frame rate between 5,000 and 20,000 was used, depending on the velocity of the water-in-oil droplets.

The mixing was evaluated by calculating the relative mixing index (RMI) defined by Hashmi&Xu^67^:

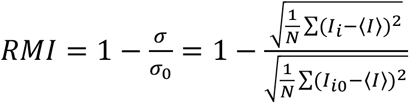

Here σ and σ_0_ are standard deviations of the intensity inside the drop at the moment of measurement and at time 0, respectively. The intensity analysis was performed in ImageJ 1.53e using the freehand selection tool, and a contour around drops at a different position along the channel was drawn. For those selected areas, the pixel intensity was measured using ImageJ.

### Visualization

Images of macromolecules and density maps were prepared using UCSF ChimeraX v1.4^68^, the 3D schematics of the setup and the microfluidic chip operation were designed using Autodesk Inventor 2022, the distributions of particle orientations were plotted using a script modified from Ref ^69^.

## Acknowledgments

We thank Prof. Henri De Greve for designing primers used to clone the *lacZ* gene. We are indebted to Prof. Andreas Manz (KIST, Saarbrüken) for helpful discussions, Prof. Benoit Schied (ULB, Brussels) for a practical introduction to microfluidics and help in setting up microfabrication laboratory, and Dr. Marcus Fislage for assistance with cryo-EM data collection. We would like to acknowledge the funding provided by Vlaams Instituut Voor Biotechnologie, Fonds Wetenschappelijk Onderzoek (Grant Nos. G0H5916N, G054617N to R.G.E.), and by the European Research Council (Grant No. 726436 to R.G.E.).

## Author Contributions

S.T. developed the microfluidic chip for trEM, established conditions for the preparation of trEM samples, collected, and processed trEM data for apoferritin and β-galactosidase. R.G.E., S.T., and R.C. designed and constructed the plunger device. S.T. and M.D. optimized the treatment of EM grids for optimal droplet spreading. M.D. collected and analyzed trEM data for GroEL:GroES. A.S. produced and purified proteins. R.G.E prepared the original manuscript draft. S.T., M.D., and R.G.E. prepared figures, reviewed, and edited the manuscript. R.G.E. conceived, managed, and supervised the project and acquired funding.

## Supplementary information

**Supplementary Fig. 1.**
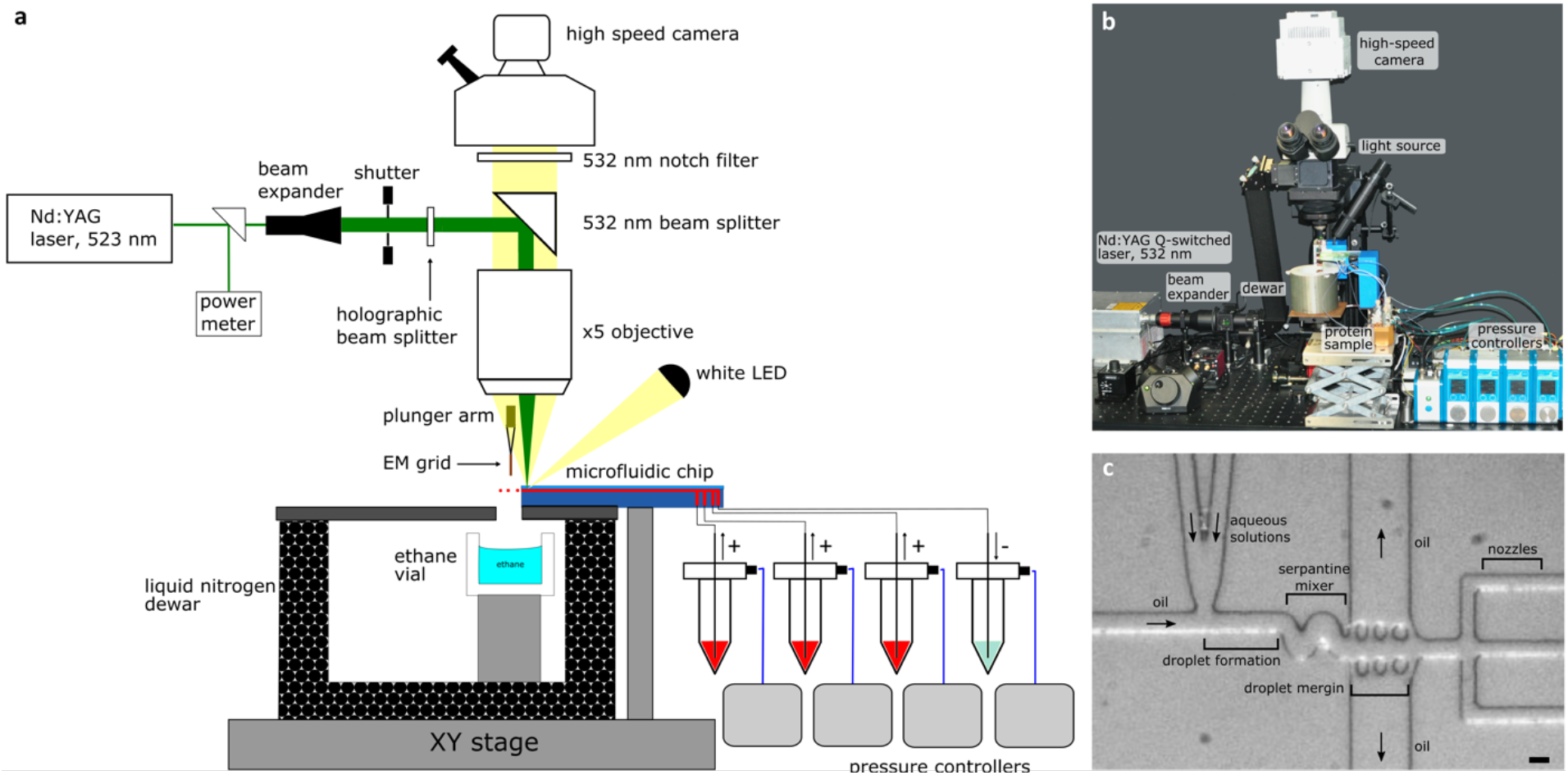
Time-resolved plunger setup. **a**, Schematics of the time-resolved device. The key elements of the setup are labeled, and the laser (green) and white light beams used for high-speed recording (yellow) are shown. **b**, Photograph of the time-resolved plunger setup.**c**, Photograph of a microfluidic chip used for performing trEM experiments. The functional modules are indicated. Scale bar 50 μm.

**Supplementary Fig. 2.**
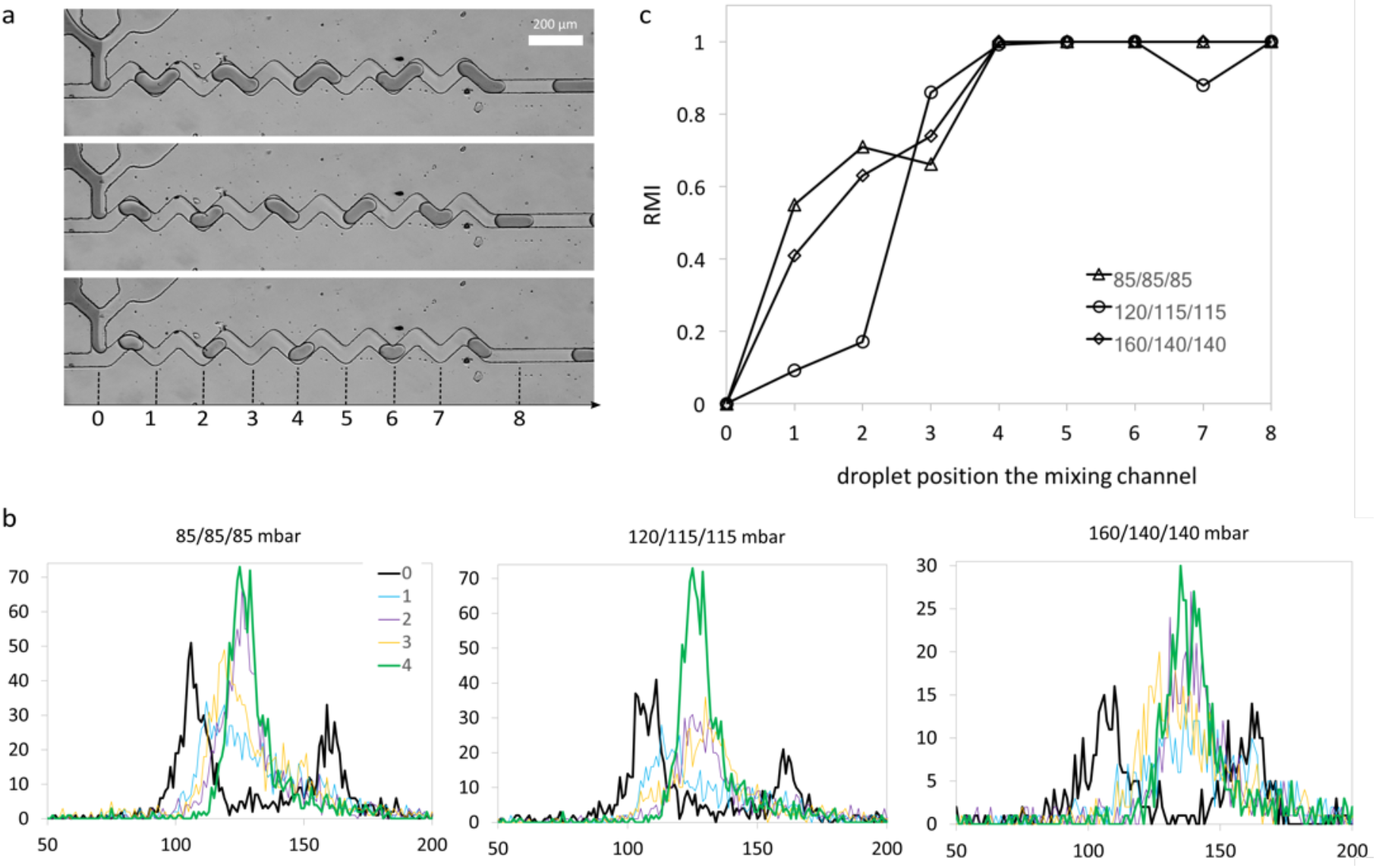
Evaluation of in-drop mixing efficiency. **a**, Experimental design: serpentine channel with 16 sections was used to evaluate mixing efficiency. Position 0 is the position at which two solutions flow parallel to each other before droplet formation. Pressures applied to the oil and water phase from top to bottom were 80, 85; 120,115; and 160, 140 mbar. Positions used for evaluating the relative mixing index (RMI) are indicated. **b**, Pixel intensity histograms of the imaged droplets qualitatively indicate the mixing extent. The histograms represent pixel intensity distributions within identical areas of the water-in-oil droplet at the indicated positions. **c**, RMI for three flow conditions calculated at eight selected positions.

**Supplementary Fig. 3.**
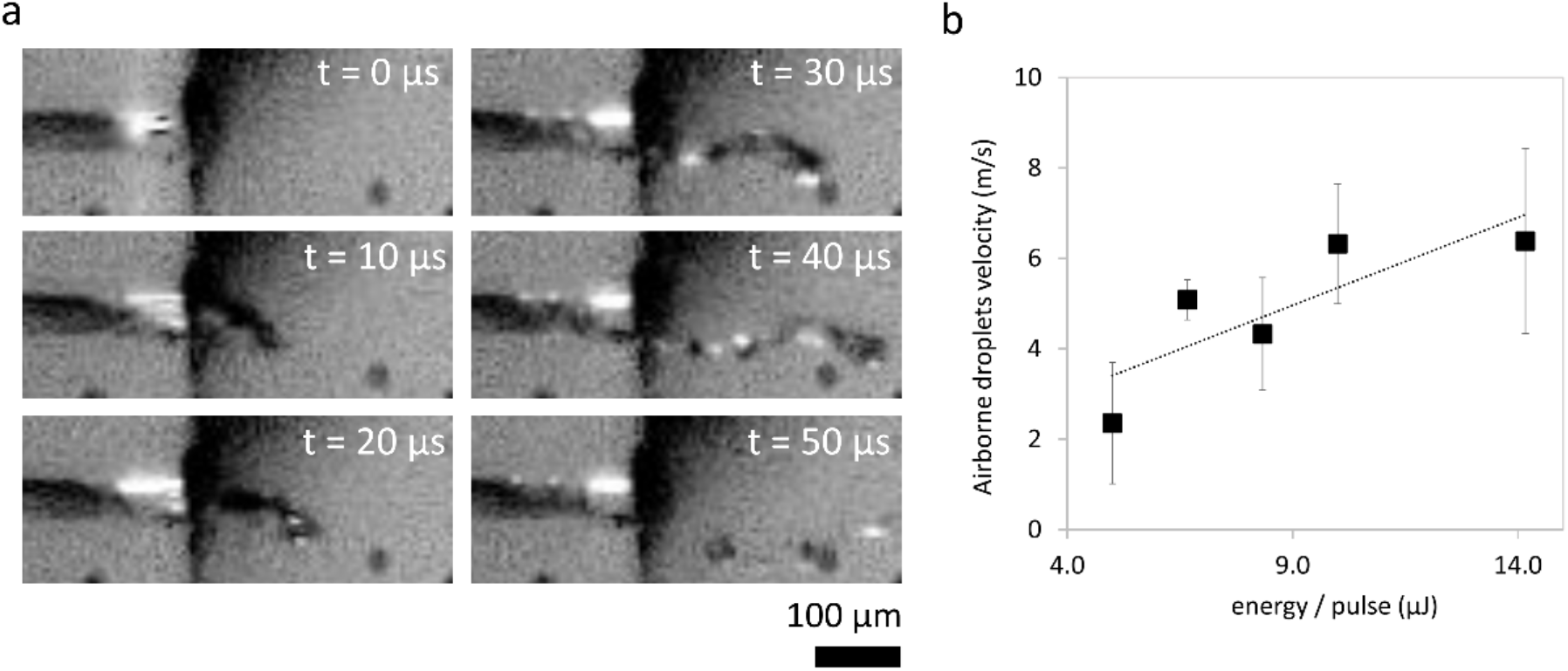
Droplet velocity in the jets generated by LIC. **a**, Snapshots of laser-induced jet extracted from a high-speed video recorded at 100,000 fps starting from the laser pulse. The images show the formation of a jet and its disintegration into droplets. **b**, Dependence of jet velocity on the laser power. The measurements were performed at a laser frequency of 3500 Hz. The energy per pulse is scaled by a factor of 12 to account for the splitting of the laser beam into 6 beams by holographic plate and power reduction by the beam splitter. The concentration of amaranth dye was 16 mM (10 mg/mL).

**Supplementary Fig. 4.**
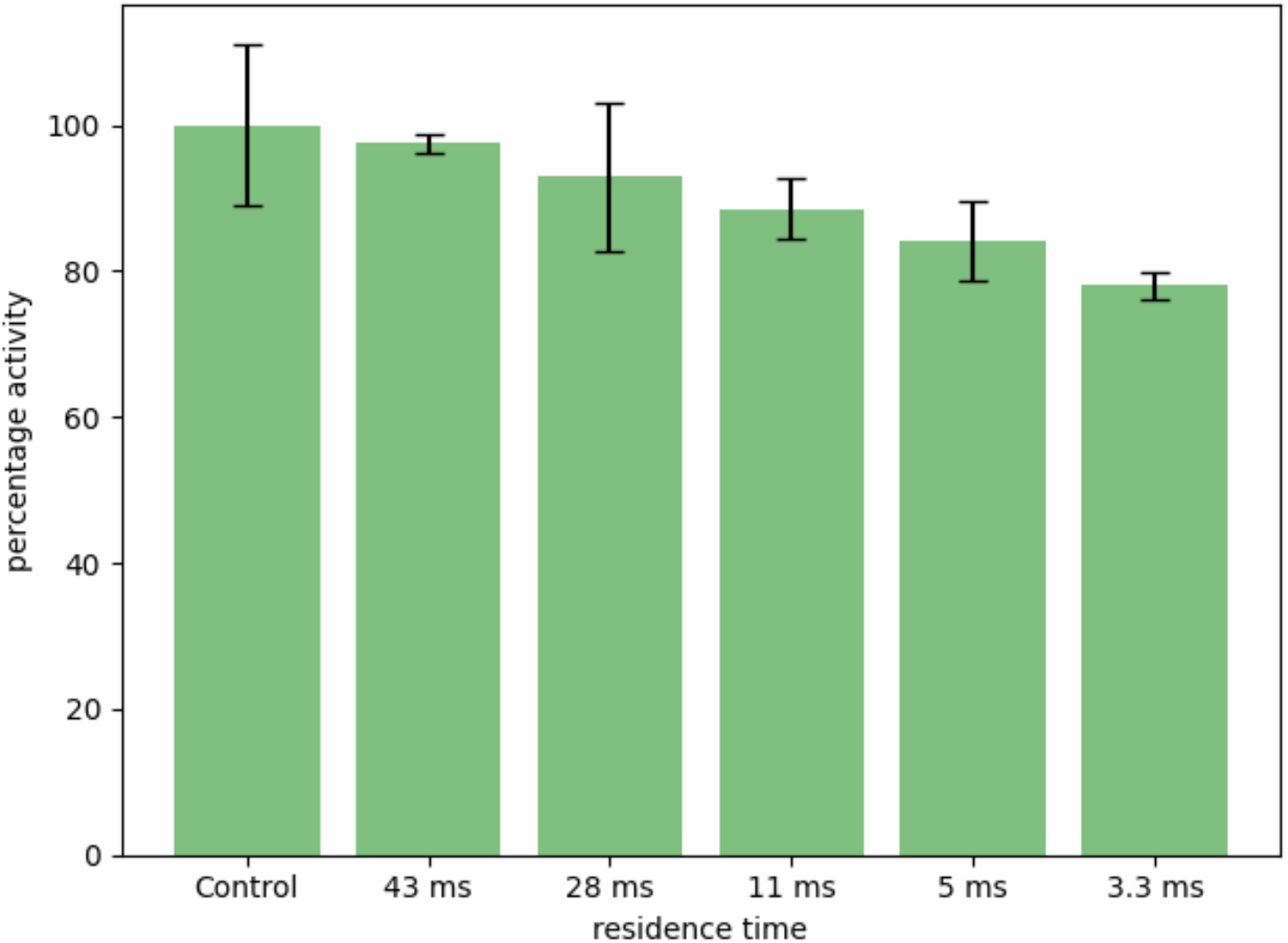
Influence of microfluidic chip and LIC on the activity of β-galactosidase. The activity of β-galactosidase was measured after passing through the microfluidic chip under conditions used for trEM plunging, that is using a similar concentration of amaranth dye, droplet-based mixer, and LIC. The sprayed solution was collected, and its enzymatic activity was measured. The control sample was not passed through the chip but contained amaranth dye in the buffer. The activity values were scaled by average control activity. All measurements were repeated 3 times.

**Supplementary Fig. 5.**
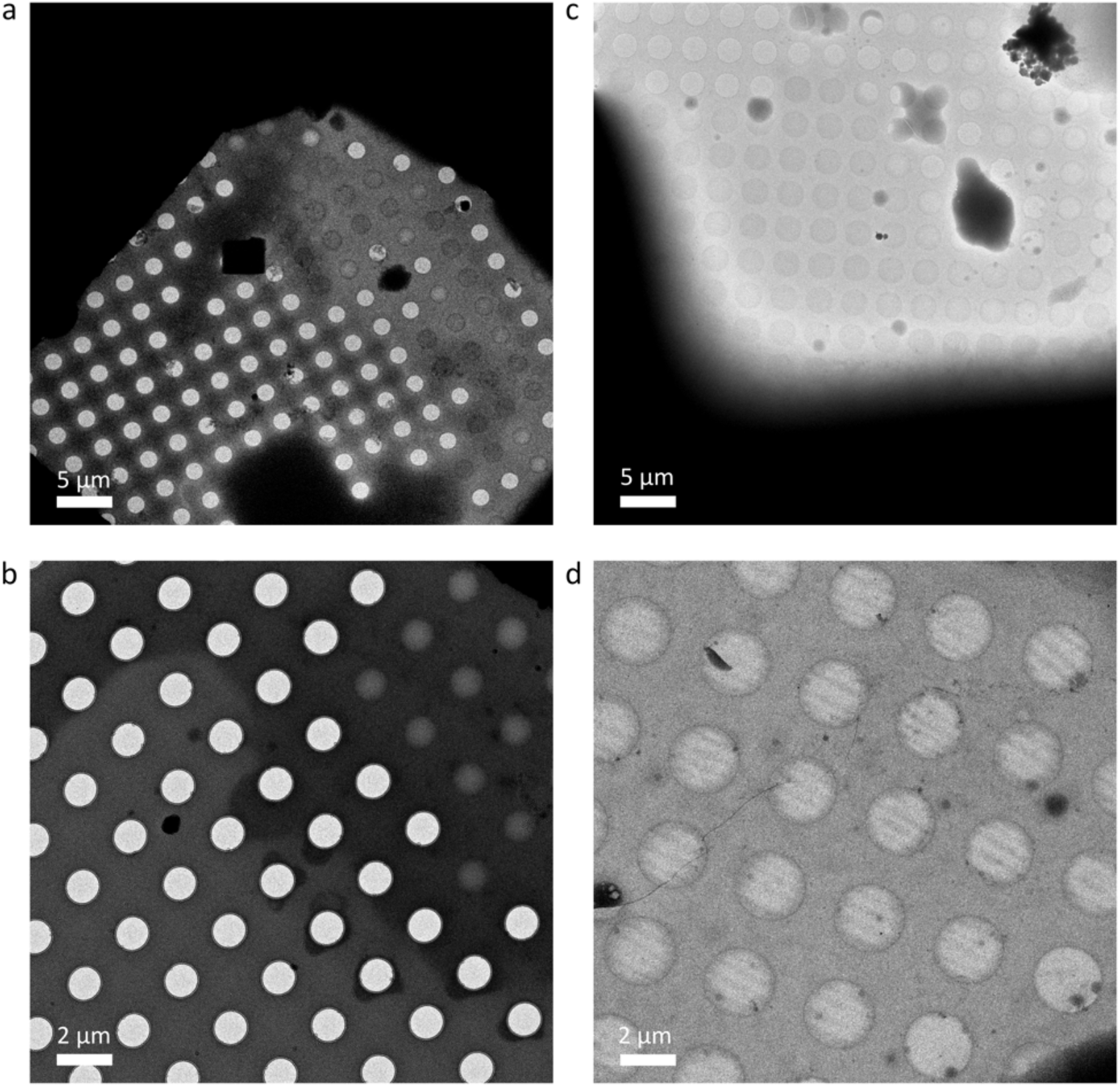
Spreading of microdroplets on holey carbon grids. **a**, **b**, Examples of spreading of LIC-generated droplets on a holey carbon grid (t_plunge_ = 100 ms) and **c, d**, on a holey carbon grid coated with 3 nm thick carbon layer (t_plunge_ = 200 ms). On holey grids, many holes remained empty even though the liquid spread over the carbon. On grids with an additional continuous carbon layer, many holes were covered with a thin layer of vitreous ice.

**Supplementary Fig. 6.**
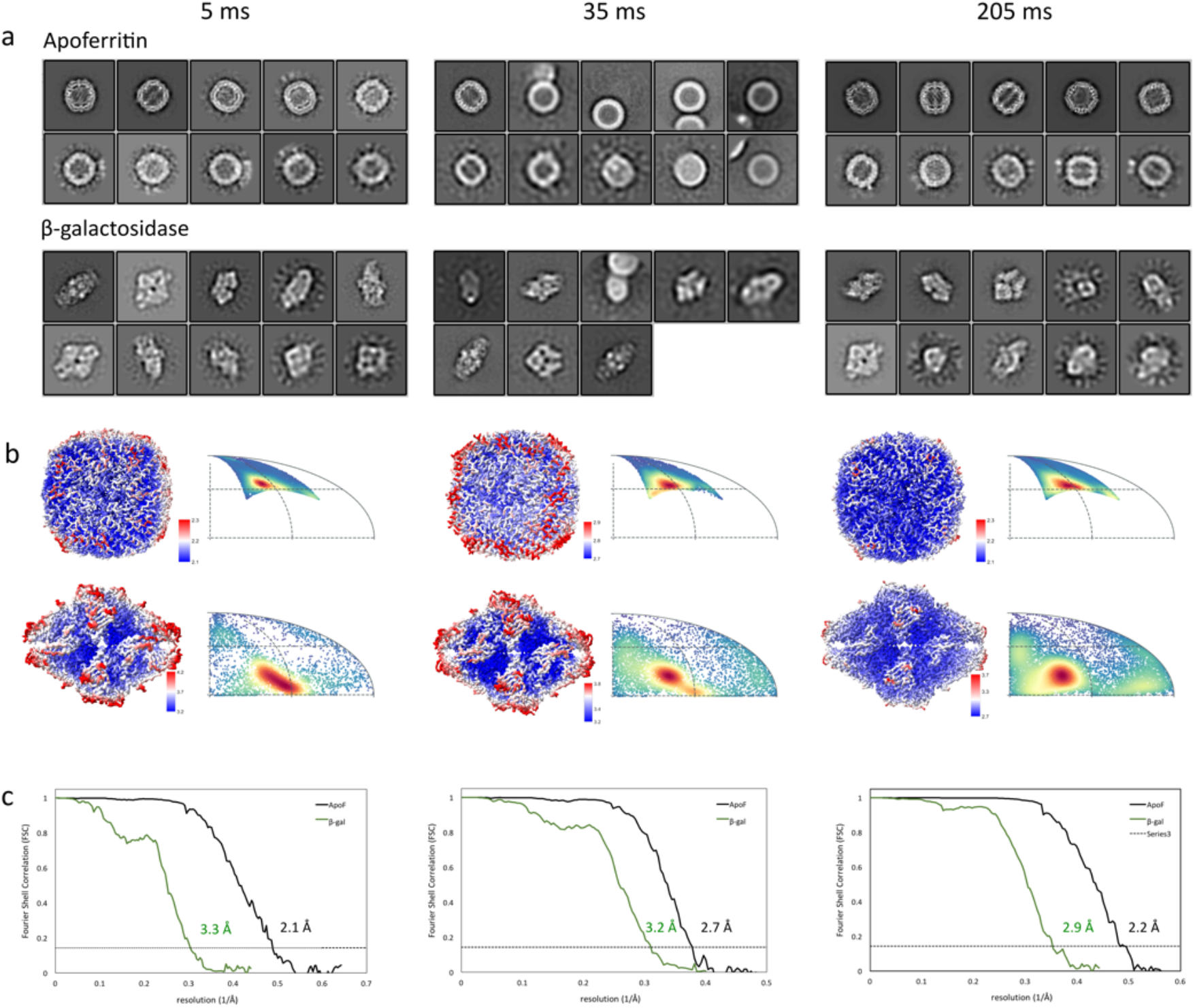
Processing of trEM data for apoferritin and β-galactosidase. **a**, 2D class averages. Classes were separated by their appearance on the classes corresponding to apoferritin and to β-galactosidase particles. **b**, Reconstructions colored according to local resolution, and distribution of particle orientations. **c**, Fourier Shell Correlation (FSC) plots for half-maps.

**Supplementary Fig. 7.**
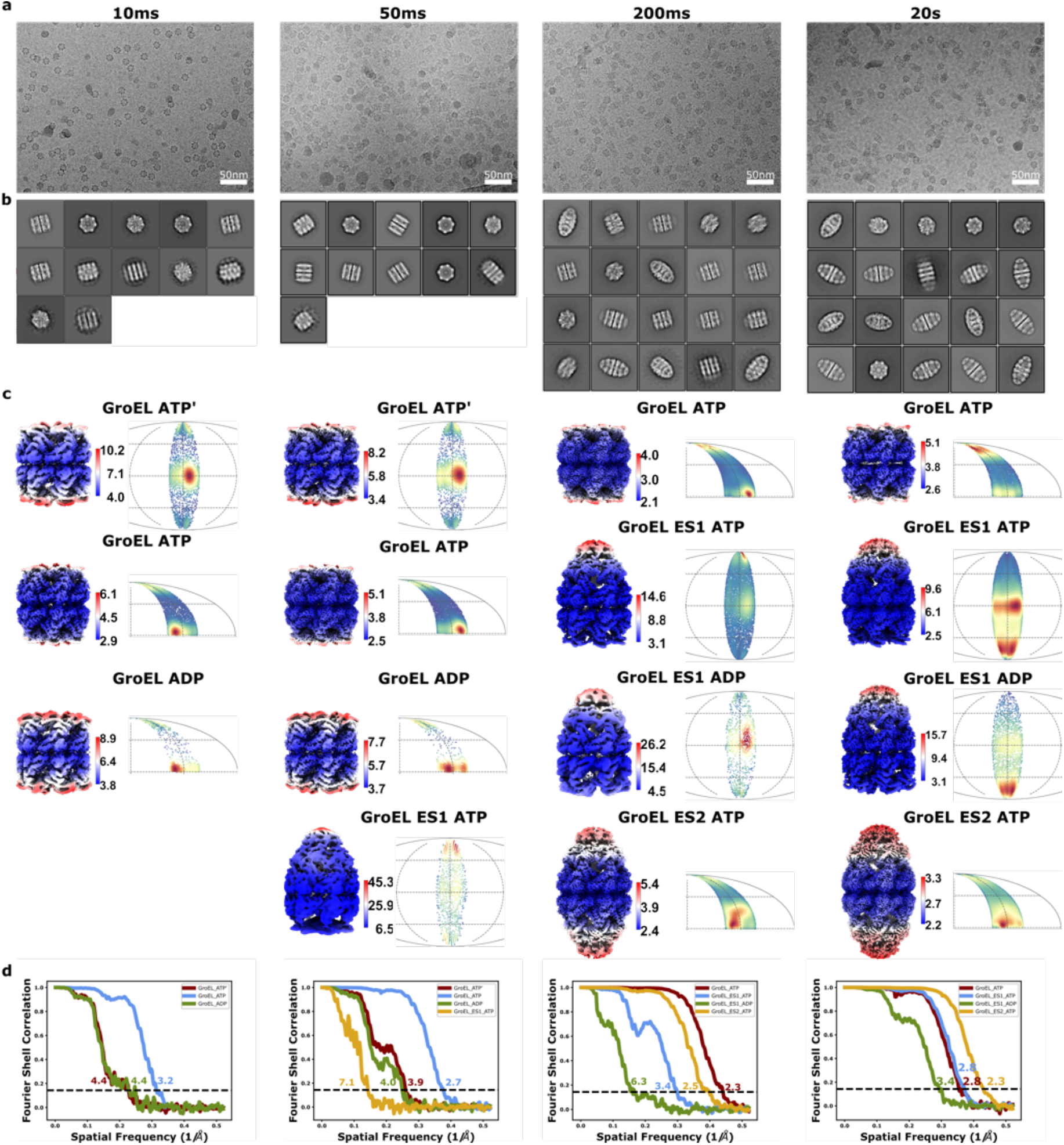
Properties of trEM data and reconstructions for GroEL:GroES data. **a**, Representative cryo-EM micrographs shown for individual reaction time points, **b**, 2D class averages. **c**, Reconstructions colored according to local resolution. The distribution of particle orientations is shown for each reconstruction. **d**, Half-maps FSC plots.

**Supplementary Fig. 8.**
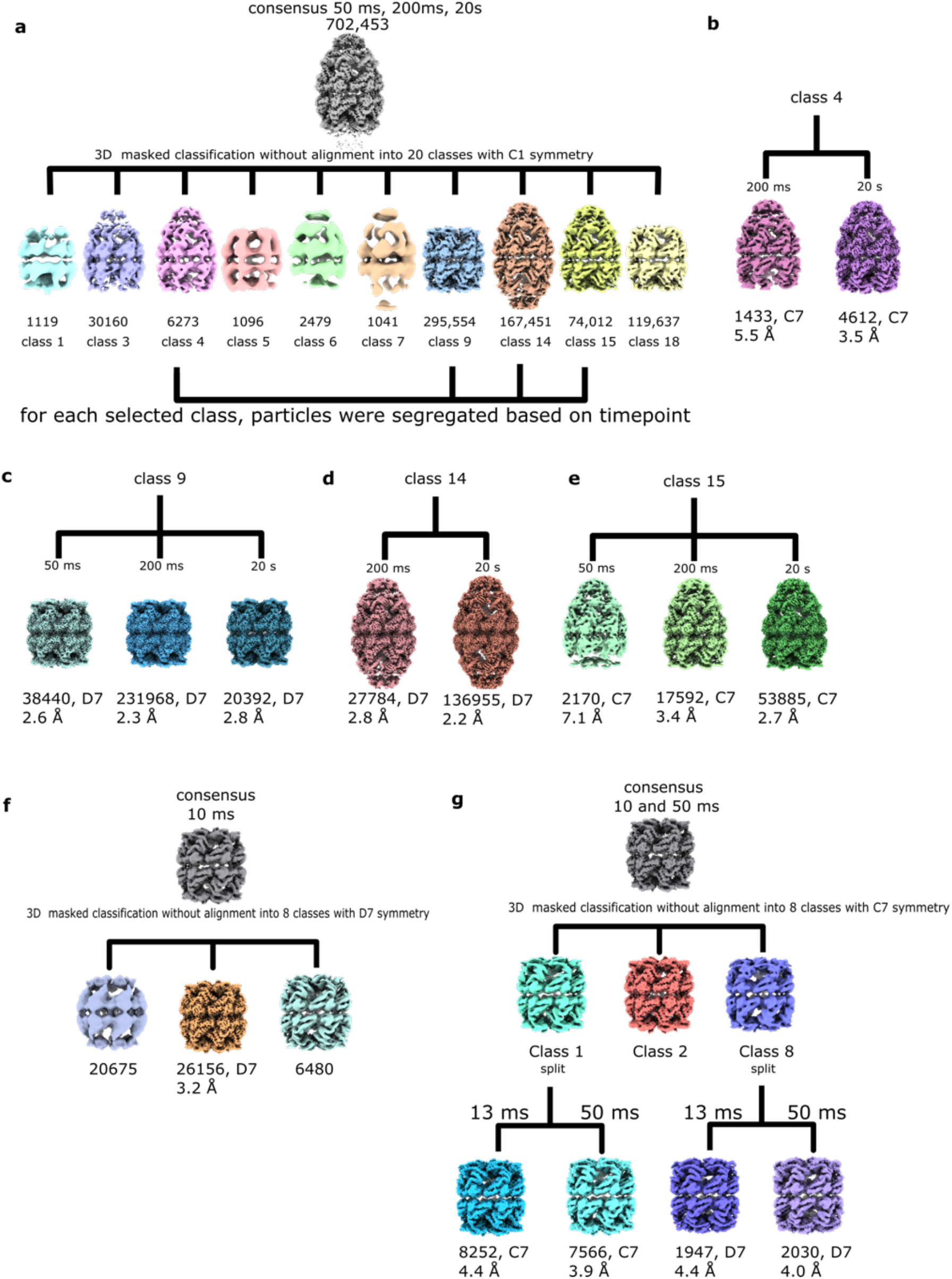
Processing scheme for trEM GroEL:GroES data. **a,** Masked 3D classification was performed after consensus 3D refinement of pooled data for time points 50, 200 ms, and 20 s. The particles were classified into 20 classes. Ten classes with a significant number of particles are shown out of which classes displaying high-resolution features were retained. Each of these classes was split into subsets corresponding to individual timepoint. **b-e,** Reconstructions from data subsets corresponding to individual time points for panel **a,** classes 4, 9, 14, and 15, respectively. **f**, Processing scheme for 13 ms timepoint. A consensus volume map was 3D classified with a mask into 6 classes. **g**, Data from 13 and 50 ms timepoints were merged, a consensus map was auto-refined with C7 symmetry, and particles were classified into 8 classes. Three classes converged to high resolution and are shown in the scheme. Class 1 and class 8 were split into 13 and 50 ms subsets to generate time point-specific individual volumes.

**Supplementary Table 1.** Cryo-EM statistics of time-resolved data collected from apoferritin mixed with β-galactosidase.

| Data collection | 5 ms |  | 35 ms |  | 200 ms |  |
| --- | --- | --- | --- | --- | --- | --- |
| Electron microscope |  |  | CryoARM300 |  |  |  |
| Electron detector |  |  | K3, CDS mode |  |  |  |
| Voltage (kV) |  |  | 300 |  |  |  |
| Nominal magnification |  |  | 60,000 |  |  |  |
| Exposure time (s) |  |  | 2.796 |  |  |  |
| Defocus range (μm) |  |  | 0.5-3.5 |  |  |  |
| Number of exposures per stage position |  |  | 5 |  |  |  |
| Number of frames |  |  | 59 |  |  |  |
| Pixel size (Å) | 0.743 |  | 0.750 |  | 0.746 |  |
| Electron dose (e- Å<sup>-2</sup>) |  |  | 59 |  |  |  |
| Number of micrographs used for particle selection | 1733 |  | 722 |  | 1409 |  |
| 3D reconstruction |  |  |  |  |  |  |
|  | AF* | β-gal* | AF | β-gal | AF | β-gal |
| Final particles | 12,593 | 4,013 | 7,432 | 6,204 | 60,146 | 12,887 |
| Applied symmetry | O | D2 | O | D2 | O | D2 |
| Resolution (Å) | 2.1 | 3.3 | 2.7 | 3.2 | 2.2 | 2.9 |
| Sharpening B-factor (Å<sup>2</sup>) | -38 | -50 | -76 | -42 | -55 | -54 |

\* AF – apoferritin, β-gal – β-galactosidase

**Supplementary Table 2a.**
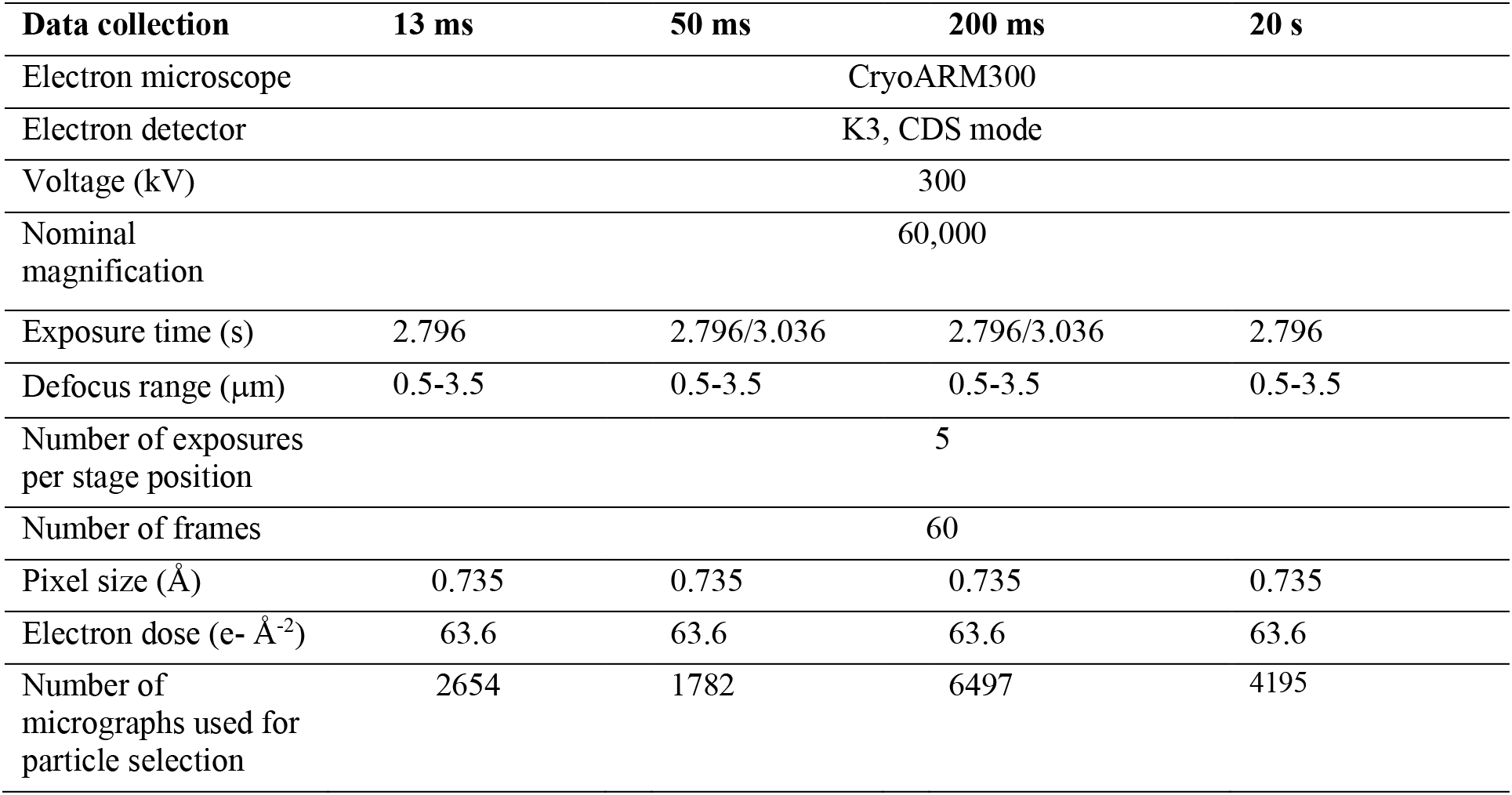
Cryo-EM statistics of time-resolved data collected from GroEL:GroES complexes.

| Data collection | 13 ms | 50 ms | 200 ms | 20 s |
| --- | --- | --- | --- | --- |
| Electron microscope | CryoARM300 |  |  |  |
| Electron detector | K3, CDS mode |  |  |  |
| Voltage (kV) | 300 |  |  |  |
| Nominal magnification | 60,000 |  |  |  |
| Exposure time (s) | 2.796 | 2.796/3.036 | 2.796/3.036 | 2.796 |
| Defocus range ($\mu\text{m}$) | 0.5-3.5 | 0.5-3.5 | 0.5-3.5 | 0.5-3.5 |
| Number of exposures per stage position | 5 |  |  |  |
| Number of frames | 60 |  |  |  |
| Pixel size ($\text{\AA}$) | 0.735 | 0.735 | 0.735 | 0.735 |
| Electron dose ($\text{e}^- \text{\AA}^{-2}$) | 63.6 | 63.6 | 63.6 | 63.6 |
| Number of micrographs used for particle selection | 2654 | 1782 | 6497 | 4195 |

**Supplementary Table 2b.** Cryo-EM statistics of time-resolved data collected from GroEL:GroES complexes: properties of the reconstructions.

| Time | 13 ms |  |  |  | 50 ms |  |  |  | 200 ms |  |  |  | 20 s |  |  |  |
| --- | --- | --- | --- | --- | --- | --- | --- | --- | --- | --- | --- | --- | --- | --- | --- | --- |
| | Resolution ($\text{\AA}$), $N_{\text{particles}}$, sharpening B-factor ($\text{\AA}^2$), symmetry | | | | Resolution ($\text{\AA}$), $N_{\text{particles}}$, sharpening B-factor ($\text{\AA}^2$), symmetry | | | | Resolution ($\text{\AA}$), $N_{\text{particles}}$, sharpening B-factor ($\text{\AA}^2$), symmetry | | | | Resolution ($\text{\AA}$), $N_{\text{particles}}$, sharpening B-factor ($\text{\AA}^2$), symmetry | | | |
| GroEL-(ADP) | 4.4 | 1947 | -106 | D7 | 4.0 | 2030 | -95 | D7 |  |  |  |  |  |  |  |  |
| GroEL-(ATP)/(ADP) | 4.4 | 8252 | -142 | C7 | 3.9 | 7566 | -106 | C7 |  |  |  |  |  |  |  |  |
| GroEL-ATP | 3.2 | 26156 | -88 | D7 | 2.7 | 38440 | -73 | D7 | 2.3 | 231968 | -70 | D7 | 2.8 | 20392 | -67 | D7 |
| GroEL-ATP:GroES |  |  |  |  | 7.1 | 2170 | -396 | C7 | 3.4 | 17592 | -86 | C7 | 2.7 | 53885 | -80 | C7 |
| GroEL-ATP/(ADP):GroES |  |  |  |  |  |  |  |  | 6.3 | 1433 | -104 | C7 | 3.4 | 4612 | -75 | C7 |
| GroEL-ATP:2GroES |  |  |  |  |  |  |  |  | 2.5 | 27784 | -64 | D7 | 2.3 | 136955 | -72 | D7 |

**Supplementary Table 3.** Settings and conditions used for plunging trEM grids.

| Proteins and actuator molecule | Reaction time (ms) | Plunge time (ms) | Final amaranth concentration (mM) | Oil / protein channel pressure (mbar) | Laser energy (μJ/pulse) | Laser frequency (kHz) |
| --- | --- | --- | --- | --- | --- | --- |
| AF/ β-gal | 5 | 3 | 16 | 360 / 180 | 80 | 4 |
| AF/ β-gal | 35 | 20 | 12 | 90 / 80 | 120 | 3 |
| AF/ β-gal | 205 | 200 | 8 | 100 / 90 | 130 | 3.5 |
| GroEL / ATP | 13 | 5 | 19 | 120-130/90 | 47 | 4 |
| GroEL / GroES / ATP | 50 | 35 | 13 | 90/70-80 | 80 | 5 |
| GroEL / GroES / ATP | 190-200 | 170-180 | 13 | 90/70-80 | 80 | 5 |

**Supplementary Video 1 High-speed imaging of functioning trEM setup**. The video is composed of 4 parts showing how microfluidic chip and plunger work. **Part 1**. Generation of microjets by LIC. The formation of airborne jets after each laser pulse is visualized using a microfluidic chip consisting of one single channel with a nozzle dimension of ≈50 × 50 μm (width x depth). **Part 2**. Operation of the complete device shows how the 2 sample solutions are encapsulated in droplets mixed, then merged in a single stream, and microjets are generated from three nozzles by LIC. The ejected droplets are applied on the EM grid moving towards the liquid ethane vial. **Part 3** shows the trajectory of the plunger arm and grid during plunging. First, the grid is slowly brought close to the nozzle where it is accelerated towards the ethane vial and decelerated to avoid the arm recoil. Time is indicated relative to the moment when the grid passes in front of the nozzles. In **part 4** the operation of the device in pulsed mode is shown.

The software controlling the device allows temporary application of the pressure to the chip channels and reduces flow in between the plungings to reduce sample consumption. In this example, the standby pressures are 36, 36, 70, and 0 mbar, whereas during plunging the flow is driven by applying pressures of 140, 140, 150, and −10 mbar over a time span of around 200 ms. The slowdown of flow is slower than depressurization likely reflecting the elastic relaxation of the chip.

